# Current Models of Speech Motor Control: A Control-Theoretic Overview of Architectures & Properties

**DOI:** 10.1101/197285

**Authors:** Benjamin Parrell, Adam C. Lammert, Gregory Ciccarelli, Thomas F. Quatieri

## Abstract

This paper reviews the current state of several formal models of speech motor control with particular focus on the low level control of the speech articulators. Further development of speech motor control models may be aided by a comparison of model attributes. The review builds an understanding of existing models from first principles, before moving into a discussion of several models, showing how each is constructed out of the same basic domain-general ideas and components – e.g., generalized feedforward, feedback, and model predictive components. This approach allows for direct comparisons to be made in terms of where the models differ, and their points of agreement. Substantial differences among models can be observed in their use of feedforward control, process of estimating system state, and method of incorporating feedback signals into control. However, many commonalities exist among the models in terms of their reliance on higher-level motor planning, use of feedback signals, lack of time-variant adaptation, and focus on kinematic aspects of control and biomechanics. Ongoing research bridging hybrid feedforward/feedback pathways with forward dynamic control, as well as feedback/internal model-based state estimation is discussed.

## I. INTRODUCTION

Several formal models of speech motor control have been formulated and presented in the speech production literature. Based on decades of observation, it seems clear that the mechanisms of speech motor control are complex, and consequently benefit from the detailed and rigorous description that formal, mathematical models can provide. Speech motor control is, indeed, one of the most intricate sensorimotor activities in which humans engage. Producing speech requires fine timing and coordination of muscles that are interwoven, redundant and have complex mechanical properties, in order to move the diverse articulatory structures of the tongue, lips, jaw, velum and larynx into a wide range of configurations, all of which have a nonlinear relationship with the vocal tract’s acoustic output. Control mechanisms are additionally modulated by higher-level processes that determine motor planning, and also mediate semantic, syntactic, prosodic and phonological organization. The various aspects of speech motor control can be conceptualized as layered modules (see Figure 1). In such a layered description, the bridge between higher-level planning processes and the movements of the biomechanical speech-producing structures is a layer which produces motor commands that drive kinematics given some motor plan and potentially in light of some monitoring or prediction of action. The central role filled by this layer – hereafter, simply referred to as the *control layer* – has ensured that all formal models of sensorimotor control for speech have defined architectures that govern its functionality. The field of models that have provided a formal description of the control layer comprises: DIVA (Guenther, 1994, 2016), Task Dynamics (Saltzman and Kelso, 1987; Saltzman and Munhall, 1989), State Feedback Control (Houde and Nagarajan, 2011), ACT (Kröger et al., 2009), GEPPETO (Perrier et al., 2005), FACTS (Ramanarayanan et al., 2016).

**FIG. 1.**
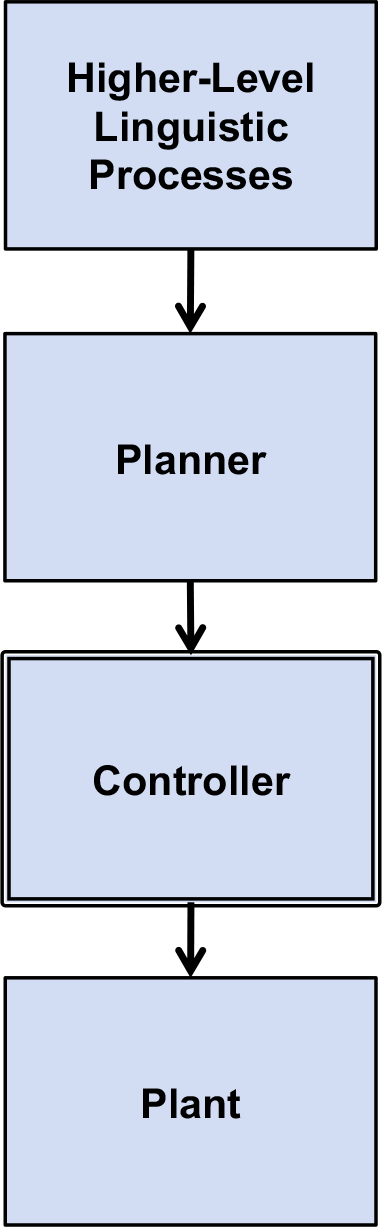
Representation of the distinct levels of speech production modeling. This paper focuses on modeling the speech controller, the system that takes in a speech plan and potentially feedback from the plant and issues motor commands to the plant. Other components of the speech production hierarchy include higher level linguistic processes (prosody, semantics, syntax), the planner (low level sequencing of motor actions), and the plant itself (e.g. speech synthesizers including but not limited to articulatory synthesizers such as CASY, Birkholz or Maeda).

An impediment to progress in developing rigorous speech motor control models appears to be the variety of distinct approaches, taken in the published literature, to explaining the attributes of the more prominent models of speech motor control. There is very little agreement, for instance, even concerning the terminology used to describe the models. Nevertheless, there is reason to believe that a direct comparison of speech control models is possible, based on the important, high-level observation that the models presented in the literature are all closely related to engineering approaches to motor control, and bear a strong resemblance to classical control-theoretic architectures. Given that the theory behind current understanding of biological motor control largely grew out of early advances in engineering fields (Bellman, 1957; Wiener, 1948), it is perhaps unsurprising that the same is true specifically in the area of speech motor control. Indeed, engineering approaches are a sensible place to begin investigations into the nature of speech motor control, in part because our current understanding of the functional interpretation of motor control neuroanatomy follows the engineering architectures closely (consider, e.g., Brainard and Doupe (2002); Shadmehr and Krakauer (2008); Takakusaki (2017); Wolpert *et al.* (1998)).

Progress in the development of speech motor control models may be facilitated by a direct comparison of the various models, using a common framework of domain-general (i.e., not speech-specific) motor control principles and unified terminology to describe their attributes. The purpose of the present paper is to provide such a direct comparison for models of the control layer that utilize mechanisms to move the plant in support of accomplishing speech tasks in accordance with higher-level speech goals. These models have been developed to attempt meaningful reproduction of speech behavior, including potentially acoustics, articulatory and neural signals. Demonstrations of the ability of these models to capture aspects of human speech production kinematics have been presented in the literature, and the extent and quality of these efforts may differ by model. No systematic review will be offered here of experimental data, either behavioral or neurological, that has been or could be used to support the expressivity or biological plausibility of any model. However, a brief summary of the demonstrated capabilities of each model is included. This choice reflects an intention to focus on the model architectures themselves.

Our review begins with general motor control principles and approaches, before moving into basic, domain-general models of motor control. The paper then proceeds to provide detailed discussions of currently proposed models of speech motor control, showing how each model is constructed out of these basic domain-general ideas and components. By showing how each model is built up on these basic elements, this approach allows for a clear comparison between the proposed models, showing where they differ as well as points of agreement. The present review focuses specifically on control of the speech articulators in fully developed, adult speech. Control that is adaptive (i.e., time variant), which may be relevant for speech acquisition and development, will only be considered in the discussion, and not in the primary overview framework. Formal explanations, including an appendix with full equations for each model, is provided where possible. Other important aspects of speech production, including learning and optimization, higher-level linguistic processing, motor program generation (i.e. the “planner”), the neurological basis of hypothesized model components, and biomechanical details of the speech articulators (i.e. the “plant”) will only be discussed to the extent necessary to clarify the nature and operation of the proposed control mechanisms.

## II. BACKGROUND

### A. Motor control principles and terminology

The first step in discussing speech motor control models is to define certain key concepts and terminology. To illustrate these ideas, a simple example is borrowed from the control of upper extremity reaching control, as shown in Figure 2, which is based on the description of a simple two-link robotic arm moving on a planar surface. This commonly-used example, though taken from a completely different domain of motor control, shares many of the same concepts and terminology with speech motor control, and has the benefit of being low-dimensional, which makes it possible to represent the relevant spaces in a two-dimensional plot. Fundamental similarities and distinctions between this simple example and the (considerably more complex) speech production system, in terms of their assumptions and structure, will be drawn where appropriate throughout the present section.

**FIG. 2.**
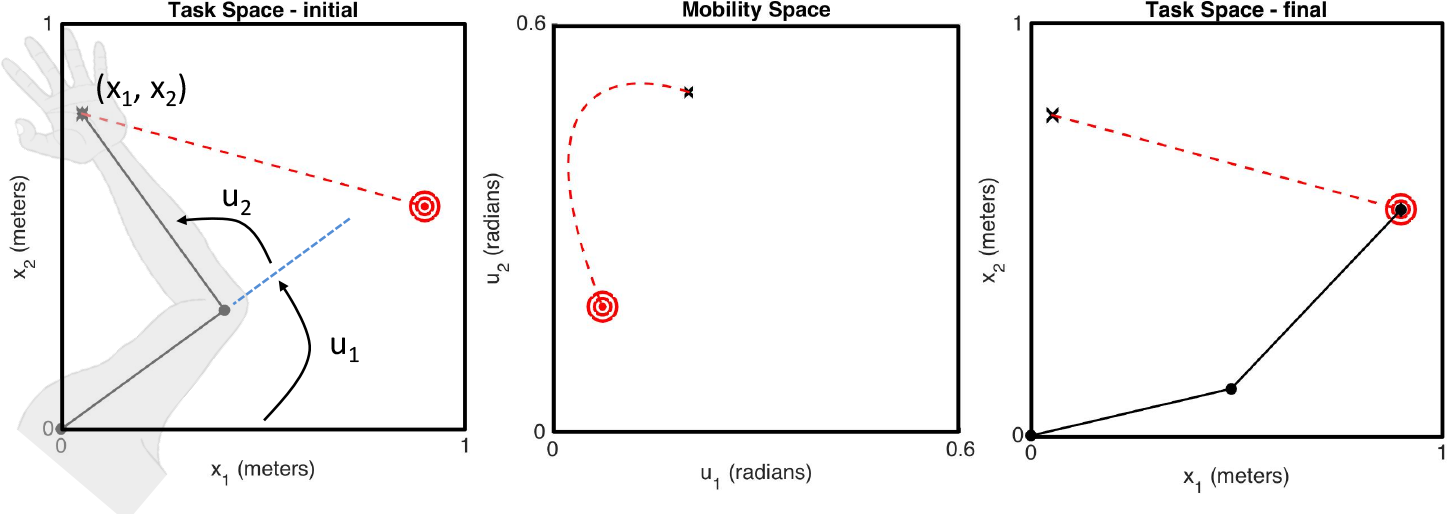
(left panel) Robot arm in its initial configuration at (*x*_1_, *x*_2_) in task space, and the final goal (red circle). The arm’s state variables (*u*_1_, *u*_2_) are defined as the angles of the shoulder and elbow. (*u*_1_, *u*_2_) are the parameters directly changed by the controller and therefore exist in mobility space. (middle panel) The trajectory in mobility space. The evolution of the mobility space variables (*u*_1_, *u*_2_) over time may be a non-linear trajectory despite a linear trajectory in task space. (right panel) The final orientation of the arm in task space at the goal.

The robotic arm, as a physical structure, is the apparatus to be controlled, and can be referred to as the plant (*G*). Note that the term *plant* is not specific to this example, and could be used in the domain of speech production to specify the vocal tract and its component articulators, as well as possibly the larynx and the respiratory system. The plant’s two links are connected to each other at a revolute joint that changes the angle between the links, *u*_2_. The proximal end of the robot’s first link is fixed at the origin of the planar surface, defined as (*x*_1_, *x*_2_) = (0, 0), but is free to rotate about this point which changes the angle *u*_1_. These two variables, *u*_1_and *u*_2_ describe the configuration of the plant, and also define the set of possible configurations of the plant, known as *mobility space*^1^. The variables *u*_1_ and *u*_2_ can be considered as elements of a single 1 − *by* − 2 vector, **u**, which can be said to specify the state of the plant in mobility space (sometimes, the *mobility state*).

The distal end of the second link (i.e., the “hand”) is considered the end-effector of the robot, the precise positioning of which is typically the focus of controlling the plant in the context of reaching tasks. The variables *x*_1_ and *x*_2_, already used to define locations on the planar surface, can also be used to describe the location of the end-effector on that surface. The space of possible locations for the end-effector is known as *task space*, and the desired outcome of a controlled movement is known as a *task*. The variables *x*_1_ and *x*_2_ can be considered as elements of a single 1 − *by* − 2 vector, **x**, specifying the state of the plant in task space (sometimes, the *task state*). Tasks with respect to the robotic arm might be putting the end-effector as a specified location in task space (i.e., achieving a state where **x** takes on a particular value), or alternatively achieving a specific trajectory through task space (i.e., tracking some sequence of values for **x**). In speech production, task spaces might include, for instance, formant space or vocal tract constriction degree/location space.

Task and mobility spaces can be viewed as “high” and “low” level spaces, respectively, with the variables comprising each space having a hierarchical arrangement where the task variables are composed of, but distinct from, mobility variables. Often this arrangement is many-to-one, such that many different (or, potentially infinite) locations in mobility space will map to the same location in task space. Task variables consequently describe the state of the plant in a way that is directly relevant to the task, and which abstracts away from a certain amount of detail as to how that task state was achieved via some mobility state. Mobility variables describe the state of the plant in a way that is more relevant to control, in the sense that motor commands are typically defined so as to affect some change in mobility state. Using the robotic arm example, motor commands would typically be given in terms of the joint angles, and not in terms of the end-effector position. In a speech context, a model might assert that motor commands are issued in terms of the positions of the speech articulators (e.g. upper lip, lower lip, tongue tip, etc.), and not in terms of some desired formant values (e.g., F1 = 500 Hz) or vocal tract constrictions (e.g., lip aperture = 2 mm).

The details of the task are specified in the *reference*, **r**, a vector representing a desired state. The reference vector typically resides in task space (**r**_**x**_), but may also be given in mobility space (**r**_**u**_) for specific applications. Reference vectors originate in the planner (*P*), and may be part of a larger motor program maintained by the planner, toward achieving some higher-level sensorimotor or cognitive goal (e.g. reach to a series of targets in space, utter the word “dad”). As implied above, however, reference vectors will typically be insufficient for use directly as motor commands to the plant because they reside in task space. The reference will need to be transformed into a motor command in mobility space. This is the function of the controller.

The controller (*C*) is the bridge between the planner and any feedback, on the one hand, and movements of the plant, on the other. The ultimate purpose of the controller is to issue motor commands that produce movement (or lack thereof) in the plant. Note that the present paper assumes that motor commands take the form of vectors in mobility space, **u**, and that those vectors can be used directly as commands to the plant. In a real biological system, several transformations may be required for encoding motor commands as neural signals, and to elicit muscle activations that bring about changes in mobility state. This assumption is made to promote consistency with the speech motor control modeling literature, and for the sake of simplicity. In any case, the motor command issued by the controller will depend either upon the reference directly, or upon the *state error*, **e**, a vector representing the difference between the reference and the plant’s state (or an estimate of that state, see below).

In biological systems, the plant’s actual state may not always be directly accessible to the controller. It can be therefore important to develop the notion of a *state estimate* (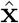 or 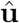), which is an internal estimate of the plant’s state, either in task space or in mobility space. The state estimate may be informed by sensory measurements of the plant’s actual state – represented by the *sensory state* vector **y** – and by predictions generated from an internal model of the plant – represented by the *predicted state* vector 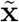 or 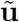. The sensory state vector, an approximation to either **x**or **u**, may be corrupted by some combination of noise (e.g., neuronal noise), delays (e.g., slowed synaptic/axonal propagation) or transformations (e.g., warping). The predicted state vector may also be imperfect, since the internal model may be inaccurate or biased. For the robotic arm example, the sensory state vector would represent measured joint angles (**y**_**u**_). This contrasts with the sensory output for speech production, which is typically considered to be a combination of auditory (**y**_*aud*_) and somatosensory signals (**y**_*somat*_), where the somatosensory signal may include proprioceptive and/or tactile sensation.

In general, motor control can be viewed as a collection of transformations between vectors and spaces of different types, and the planner, the controller, and the plant can all be described – using the conventions developed above – as functional transformations from specific inputs to specific outputs. The planner generates the reference vector, **r** = *P*(*α*) as a function of some high-level motor program-related information *α*, and possibly as a function of time: **r** = *P*(*α, t*). The controller takes a reference vector or an error vector as input and generates a motor command in mobility space: 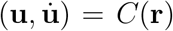 or 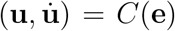. The plant, which can also be viewed as a transformation, converts motor commands, through movement, into different plant states which can be measured in both mobility and task space: 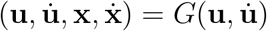. These variables are used in Figure 4, and in related diagrams throughout the paper. The state of the plant can then be measured by some sensory system: 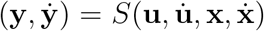, the details of which are often not explicitly treated in the literature. Therefore, the present review will often lump *G* and *S* together into a single component.

### B. Types of motor control models

The purpose of this section is to lay out, in a general way, some common control architectures that are employed in various control applications, including both controlling robotic systems as well as describing the functional aspects of physiological control. These general architectures are presented as a scaffold for understanding the specific architectures used in various speech motor control models, and also for the purpose of clarifying the terms used in the present paper to refer to those architectures. To illustrate these various architectures in an intuitive way, the example of the planar robotic arm will continue to be employed as in the previous section. However, these same architectures can be used to control much more complex systems, such as the speech production system.

#### 1. Feedback control

Figure 4b shows an example of a *feedback* system architecture that, by definition of the term, makes use of outputs from the plant for maintaining control. These feedback signals, which convey the sensory state vector, are compared with the reference vector from the planner in order to generate an error vector. The error vector, in its most basic form, simply represents the difference between the current state and the reference. The error vector is passed to the controller for determining the motor command. This type of controller is often referred to as a *closed-loop controller* in the control theory literature, since the flow of signals through the system form a loop from the motor command to the error signal and back again. Many types of controllers exist which match this general description, only a few of which will be discussed here. What all feedback controllers share is the basic idea that the error between the state of the plant (or an estimate thereof) and the reference forms the basis for the motor command issued to the plant. The simplest feedback controller design is the proportional controller, in which the motor command is simply proportional to the error signal – e.g., *C*(**e**_**x**_) = **K**_*p*_**e**_**x**_, where the term **K**_*p*_ is a matrix of weights known as the *gains*. Larger gains lead to larger motor commands (i.e. the error has more of an effect on the system) while smaller gains result in smaller commands. Smaller gains are often preferable as large gains can lead to instability and oscillatory behavior.

The second row in Figure 3 shows, across times *t*_1_, *t*_2_ and *t*_3_, the progress of the robotic arm as controlled by a feedback controller. At the beginning, the task is defined as a desired point in task space **x** = (*x*_1_, *x*_2_). This type of task is sometimes referred to as a point-attractor, or a target, since the system should evolve to approach this point in task space regardless of its initial position, given sensible motor commands that reduce the error signal over time. The motor commands issued at each time step are a function of the error, **e**_*x*_, between the current position of the end-effector and the point target. The error is determined by sensory feedback, which provides monitoring of the current state of the arm with respect to the position of the target.^2^ Although the error signal is in task space, the motor command issued by the controller must be given in mobility space since the only way to change the position of the end effector is to change the joint angles **u** = (*u*_1_, *u*_2_). The process of determining those commands requires some kind of transformation (i.e., kinematic inversion) from the desired coordinates in task space to corresponding coordinates in mobility space. Alternatively, it is also possible for the target to be a pre-specified trajectory rather than a point in task space. In this case, the error would be computed between the current position of the end-effector and the current desired position along the trajectory (typically time-locked).

**FIG. 3.**
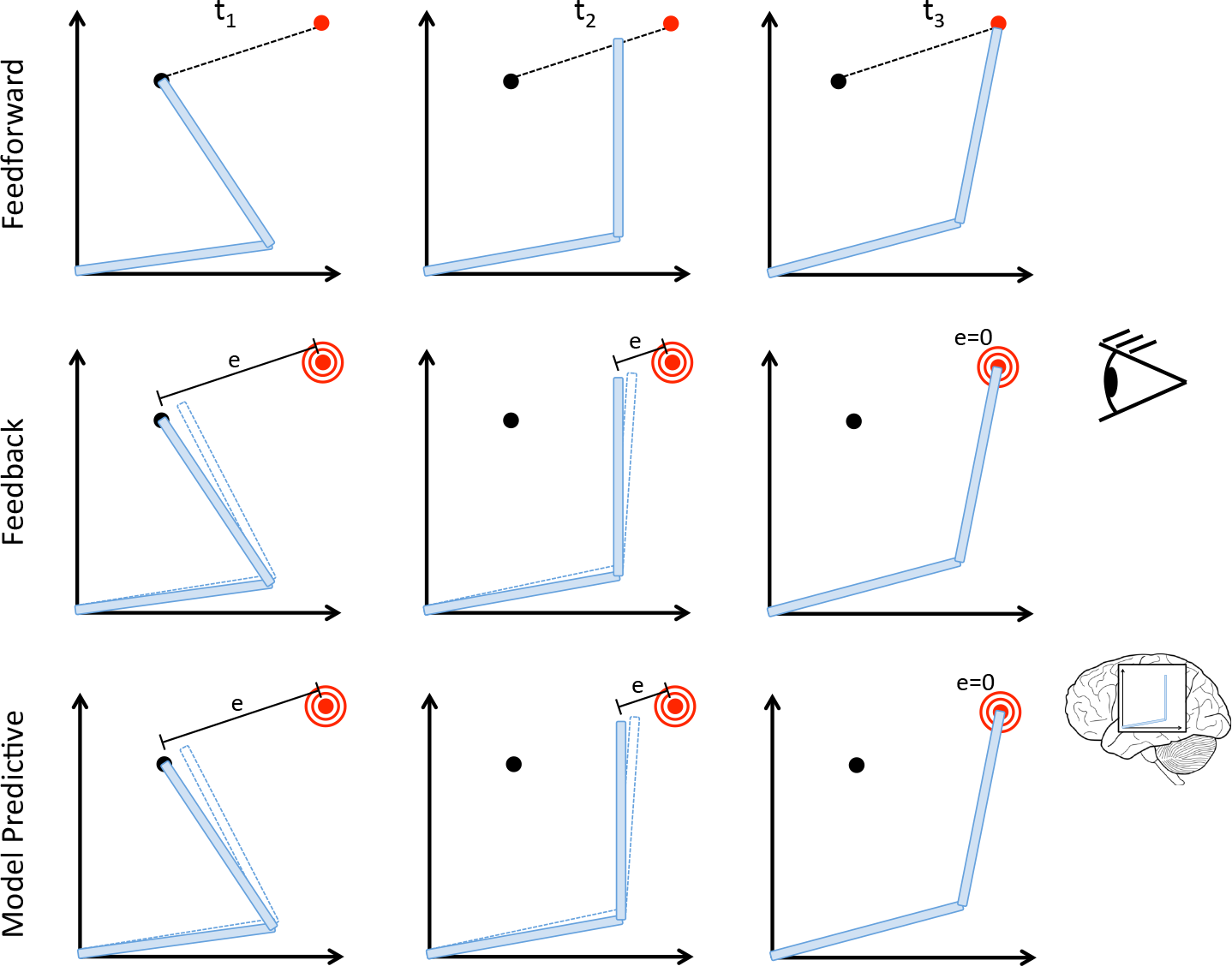
Illustration of the difference between feedforward (top row), feedback (middle row) and model predictive (bottom row) control using a simple reaching example. In feedforward control, the arm traces out a fully preplanned trajectory with no feedback about the position of the arm at any point in time. In feedback control, an error is computed between an observed state of the system (observation represented by the eye) and the target. The arm progressively works to minimize this error which drives the end effector towards the target. In model predictive control, an error is computed internally as opposed to being derived from feedback of the state of the system (represented by the brain with an internal model of the robot arm). The arm’s position is updated to minimize the predicted error of the system.

Feedback control architectures have wide applicability in engineered and biological systems. Even simple designs typically lead to systems that accurately produce desired behaviors, and which can naturally handle unstable or unpredictable environments, including external perturbations to the plant. However, feedback systems can require careful calibration to ensure stability of control. Incorrectly tuned feedback systems can result in movements that grow uncontrollably or oscillate indefinitely. Feedback architectures are also heavily dependent on the quality of feedback signals. If those signals are slow to propagate, or if they require extensive processing once received, this can lead to motor commands being issued based on outdated state information, resulting in poor and/or slow performance. Additionally, if feedback signals are corrupted or otherwise inaccurate, this can lead to inaccurate movements. These final considerations are particularly important for biological systems, as there are substantial delays and noise inherent to neural processing of sensory feedback.

#### 2. Feedforward control

One way to avoid the problems of delayed and noisy sensory information is to cut out the use of feedback entirely. Figure 4a shows an example of a system architecture that makes no use of any outputs from the plant for maintaining control. Rather, the signals issued in the system are entirely *feedforward*, with the motor commands depending only on the reference signal. This architecture is commonly referred to as an *open-loop* control system, although the terms *feedforward* and *open-loop* will be used interchangeably in the present paper. The term feedforward control is sometimes used more specifically to refer to control architectures that can monitor perturbations to the plant, and adjust the motor commands to compensate without the need for explicitly monitoring outputs from the plant, usually by employing a highly accurate mathematical model of the plant (see the section on model predictive control, below). To date, the authors are aware of only one modeling effort in the domain of speech motor control to utilize this kind of architecture (Baraduc *et al.*, 2017), with preliminary results presented.

**Fig. 4.**
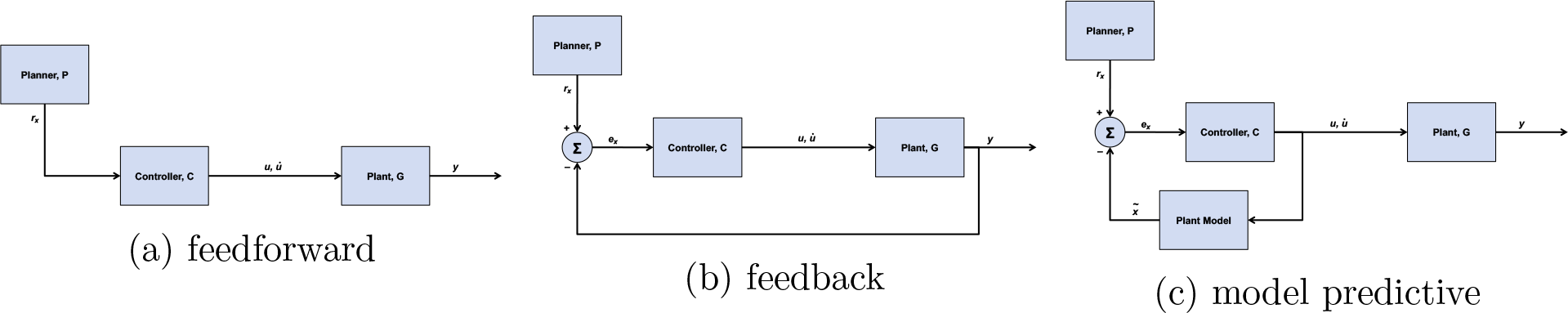
Control architecture of a generic (a) feedforward, (b) feedback, and (c) model predictive controller. The feedforward control architecture is distinguished from the other two because the controller only receives information from the planner, not information from the plant or predicted information from the plant. The feedback control and model predictive control architectures differ in the nature of the feedback received by the controller. In feedback control, the state of the plant (different than the output) is sent back to the controller. By contrast, in model predictive control, the state of the plant is sent back to the controller using an estimate of the plant based on a copy of the issued control signal.

The first row in Figure 3 shows the progress of the robot arm as controlled by a feedforward controller. From the beginning, the trajectory of the end-effector is defined in task space as a straight line originating at the end-effector’s current position. The motor commands issued to the arm at each time step are directly determined by this pre-specified trajectory. As in a feedback controller, the reference signal is defined in task space but motor commands must be issued in mobility space. Again, this requires some kind of transformation from the desired coordinates in task space to corresponding coordinates in mobility space. Although the trajectory in this example is specified in task space, as is often done, an alternative feedforward controller could define the plan in mobility space (that is, for our robot example, in terms of joint angles) or even simultaneously in mobility and task space. In any case, a key aspect of feedforward control is that no estimate of the state (that is, the arm’s estimated position) is used by the controller at any point throughout its movement. In the absence of feedback, the simplest method of generating reasonable control signals is simply to have the plan pre-specify the entire trajectory in task or mobility space, and then issue motor commands that attempt to carry out that plan step-by-step from beginning to end.

Feedforward control architectures are unsuitable for unstable or unpredictable environments, where the plant can be perturbed by interference external to the system. Without the ability to detect and monitor errors in the system output, errors tend to persist, or even compound over time. Despite this obvious disadvantage, feedforward architectures are sometime attractive because they are capable of issuing motor commands quickly and without the need for complex handling of feedback signals.

#### 3. Model predictive control

An alternative to feedforward and feedback control is *model predictive* control. A model predictive controller, like the feedforward controller, makes no use of outputs from the plant for maintaining control. However, this architecture does make use of an *internal model* of the plant, which takes motor commands as input and transforms them into a prediction of the system’s subsequent state, to predict the effects of the issued motor command. This effectively replaces feedback from the plant with a prediction of what the controller thinks that the feedback should be (Garcia *et al.*, 1989; Miall and Wolpert, 1996). An example of this architecture is shown in Figure 4c. This state prediction acts as a kind of pseudo-feedback which can be compared against the reference, producing an error signal that is provided to the controller.

Note that model predictive control can be viewed as a special case of feedforward control, if the plant model is considered to be part of the controller. This special case has been separated out as a distinct architecture in the present framework because it is central to several models of speech motor control. Therefore, feedforward architectures, as discussed here, will specifically discount architectures that are model predictive.

The third row in Figure 3 shows the progress of the robot arm as controlled by a model predictive controller. The functioning of such a controller is similar to the feedback controller example, above, in that the target is defined as desired point in task space, and the motor commands issued are a function of the error, **e**_**x**_, between the current position of the end-effector and the point target. The difference is that the error is determined by comparing the desired state to the output of an internal model.

In terms of performance, the primary advantage of such an architecture is speed, since the delays associated with predicting the plant’s state can often be much shorter than those associated with feedback propagation. Additionally, a model predictive controller is one way to avoid the need for having an entire trajectory formulated before movement begins, as is often the case with feedforward architectures. Rather, plans can be more compact, such as a single, time invariant point in task or mobility space (this is the same type of plan often used in feedback controllers). The disadvantage of these systems is that accurate internal models can be difficult to design or learn, especially for complex, nonlinear plants such as the vocal tract. A poor internal model would mean that the predicted state may not match the true state of the plant, which can result in inaccurate control. Even small errors in the prediction will accumulate over time, since there is no way of correcting the prediction. Additionally, model predictive controllers have similar problems as feedforward control architectures in dealing with unpredictable environments and perturbations.

#### 4. Combining feedforward and feedback controllers

Each basic type of control system, feedback and feedforward control, has its own strengths and weaknesses. Feedback control is stable in the face of external perturbations, but becomes inaccurate or slow when sensory information is noisy or delayed (respectively), as in most biological systems. Feedforward control can be accomplished quickly, but is unstable when the state of the system cannot be predicted due to external perturbations.

It is possible to combine some of the strengths of feedforward and feedback systems, and mitigate the weaknesses of each, by constructing a *hybrid feedforward/feedback* controller, as shown in Figure 5a. This hybrid architecture comprises separate feedforward and feedback pathways that each issue their own motor commands, a (potentially weighted) combination of which becomes the final motor command that is issued to the plant. Such an architecture has the speed of a feedforward controller, but remains sensitive to unexpected perturbations and accumulating errors. Typically, the presence of the feedforward pathway allows for lower gains to be utilized in the feedback controller, leading to better stability. The primary disadvantage of combining feedforward and feedback pathways into a single system is the introduction of more complex designs. Complex designs may be more difficult to maintain, and allow the potential for unnecessary or underutilized components. For instance, if output from the plant always equals the reference (e.g., if the environment is entirely predictable), then the feedback pathway is not utilized and essentially unnecessary, since the feedforward pathway would be sufficient for control by itself.

**FIG. 5.**
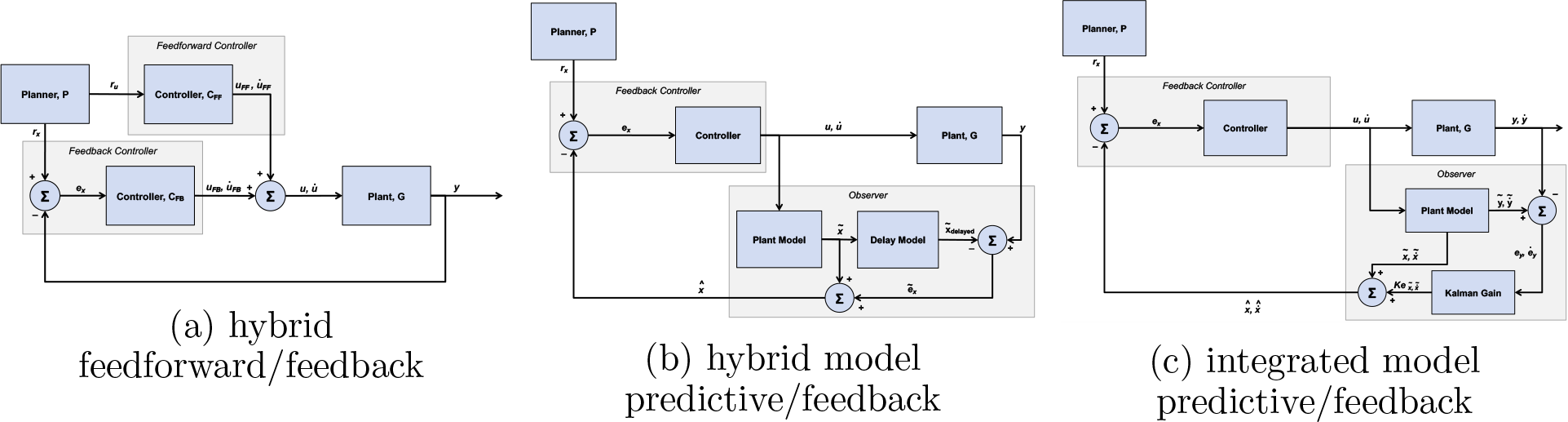
Control architectures of generic hybrid controllers. A hybrid controller uses two of the three simple control architectures discussed in Figure 4. Diagrammed here are (a) a feedforward-feedback hybrid, (b) a model predictive-feedback hybrid with simple summation of state predictions (i.e., a Smith predictor) and (c) a model predictive-feedback hybrid with full integration of state predictions (i.e., a Kalman filter). Architectures (b) and (c) are distinguished by the specific way in which model predictions and feedback are combined. In (b), the current state is estimated through a three-part error comparison processes. Architecture (c) also uses a three-part comparison, but also incorporates an observation model that maps the model prediction into sensory space, and a gain that allows for potentially variable weighting of model predictions and sensory measurement error.

One of the most useful applications of model predictive control is as a component of larger, hybrid architectures. For instance, internal model predictions can provide quick pseudo-feedback that can be used in conjunction with true feedback to provide fast, reliable control even in the face of long feedback propagation delays. Such methods are more stable than true model predictive control, since internal predictions do not need to be perfectly accurate, and small deviations between the predicted and actual states of the plant can be corrected via the feedback signal. An example of an architecture that exemplifies this concept is the Smith predictor (Ghosh, 2005; Smith, 1959), as shown in Figure 5b. A Smith predictor effectively has three error comparison processes, generating state errors serially through comparing the state with a delayed version of the internal model prediction, which in turn is compared to a non-delayed internal model prediction, with this final comparison being subsequently compared against the desired state from the reference signal. The integrated mechanisms involved in combining model predictions with feedback signals are sometimes referred to in the literature as the “observer”. The present view adopts this terminology. Note that the observer and speaker, in this conceptualization, are the same individual, as speakers observe their own speech.

A Smith predictor is not the only controller that uses both state predictions from an internal model and feedback signals. Prominent alternative approaches also use a three-part, cascaded error comparison process, but incorporate (a) an observation model, that maps the model prediction into sensory space for direct comparison with sensory measurements, and (b) a gain that allows for potentially variable weighting of model predictions and sensory measurement error. These additional aspects can afford more accurate estimation of the plant’s current state. This is the approach taken by such classic control designs as the Kalman filter (Kalman *et al.*, 1960) (Figure 5c), which provides an optimal^3^ state estimate with noisy feedback under certain strict assumptions. Importantly, the estimated state that results from combining internal predictions and feedback can be compared with the desired state to generate a motor command (Todorov, 2004), just as in a pure feedback controller. This type of controller is sometimes referred to as *state feedback control*.

### C. Speech models

The present discussion will now move from domain-general motor control theory to models of speech motor control. Among the speech production models presented in the literature, perhaps the two most prominent are DIVA (Directions Into Velocities of Articulators) and the Task Dynamics model. The development of DIVA has been driven since the mid-1990’s (Guenther, 1994) primarily by a team of researchers at Boston University, led by Frank Guenther. Task Dynamics has been developed by researchers associated with Haskins Laboratories, with Elliot Saltzman playing a key role, and with the theoretical groundwork being laid about five years prior to DIVA (Saltzman and Kelso, 1987; Saltzman and Munhall, 1989). More recent models include State Feedback Control (Houde and Nagarajan, 2011), the Feedback Aware Control of Tasks in Speech (FACTS) model (Parrell *et al.*, 2006), ACT (Kröger *et al.*, 2009), and GEPPETO (Perrier *et al.*, 2005).

Any model of speech production control must include, at a basic level, the ability to generate motor commands based on some motor plan. Those motor commands in turn activate a vocal tract model, possibly resulting in the generation of an acoustic signal. While complete models of speech production also need to include the formulation of motor plans, these elements are beyond the scope of the present paper, which focuses more narrowly on controlling the vocal tract for speech. An important reason for limiting the scope of the present paper is that the longstanding debate over acoustic vs. articulatory targets of speech production tasks is often intertwined with the critical issue of how the vocal tract is controlled. For example, DIVA’s tasks are formulated primarily in acoustic space, whereas applications of Task Dynamics (e.g., the Articulatory Phonology of (Browman and Goldstein, 1986) often assume tasks to be constrictions in the vocal tract. The choice of task space, however, is almost completely independent of the control formulations that are the focus of the current paper, and it is generally possible to reformulate any given control architecture using different task spaces. Therefore, the present work will discuss the task space used for each model, as the specific choice of task variables comprising the task spaces does differ between models, but will make no attempt to discuss the relative merits of the different task spaces used in different models. The concept of a task space is general enough to sit over and above the specific choice of task variables, while being well-defined enough as a concept to allowing meaningful comparisons of the control architectures underlying task space control.

Control elements that are relevant to any model of speech motor control, and which will be discussed in depth for each model in the following section, include: (a) the nature of feedforward mechanisms of control, including the formulation of the planner, (b) the nature and importance of feedback signals, (c) modeling of potentially imperfect sensory systems and/or perceptual processing of feedback, (d) the consequences of delays in feedforward and feedback pathways (e) the potential role of forward models in state prediction, and (f) the potential integration of both feedback and state predictions for state estimation, (g) the implementation of transformations between task space, mobility space, and sensory space, (h) the design of the controller for generating and issuing motor commands to the plant.

It is noted here that most current speech models are examples of purely kinematic controllers. That is, they do not account for dynamics or biomechanical considerations of the vocal tract. It is typically assumed that inertial parameters, centrifugal/coriolis forces and stationary external forces like gravity can all be ignored for the purposes of controller design and forward modeling. This may owe to the fact that several prominent models of the plant are purely kinematic: for instance, Maeda’s model (Maeda, 1982) and the Haskins Configurable Articulatory Synthesizer (CASY) (Iskarous *et al.*, 2003; Rubin *et al.*, 1981, 1996). The focus on kinematics may also reflect an implicit assumption that dynamics of the plant can be ignored in the domain of speech motor control. Such an assumption is quite common in robotics and human motor control, and amounts to conceptualizing the plant as a collection of stif articulators, akin to an industrial robotic arm. However, there is evidence that biomechanical factors play non-negligible roles in speech motor control (Buchaillard *et al.*, 2009; Derrick *et al.*, 2015; Nazari *et al.*, 2011; Ostry *et al.*, 1996; Perrier *et al.*, 2003; San-guineti *et al.*, 1998; Shiller *et al.*, 2002), and more recent vocal tract models such as Artisynth (Lloyd *et al.*, 2012) incorporate dynamic and biomechanical aspects in their design.

## III. PROMINENT MODELS OF SPEECH PRODUCTION

In the following section, each of the current models of speech motor control will be discussed in turn, explaining the architecture of the control system as it relates to the simple, domain-general systems discussed previously. Where necessary, additional components of each model will be touched upon, such as motor program generation. How each model addresses the key control elements listed above will also be discussed.

### A. DIVA

The Directions Into Velocities of Articulators (DIVA) model is a hybrid control system combining a model-predictive controller with separate auditory and somatosensory feedback controller loops (Golfinopoulos *et al.*, 2011; Guenther *et al.*, 2006; Tourville and Guenther, 2011). Being arguably the most complete computational model of speech motor control, DIVA has been developed to address a number of theoretical issues, primarily focused around replicating human speech production at behavioral, neurological, and developmental levels. The use of both model predictive and feedback control in DIVA is conceptually similar to a Smith Predictor. However, while a Smith Predictor uses serial error calculations to issue a single motor command, DIVA generates independent errors from each controller simultaneously. Each error is then individually transformed into a separate motor command. These three commands are then combined into a single motor command which is passed to the plant. The plant in DIVA has historically been Maeda’s model (Tourville and Guenther, 2011), but this has recently been replaced with a custom plant model (Guenther, 2016).

The basic component of the planning process in DIVA is the “speech sound”, which can be a phoneme, syllable, or multisyllabic chunk. Each speech sound is linked to three distinct tasks, each a function of time: an articulatory trajectory (often called “motor” trajectory in the DIVA literature) defined in mobility space **r**_*u*_(*t*), an auditory sensory trajectory **r**_*aud*_(*t*), and a somatosensory trajectory **r**_*somat*_(*t*). The “speech sound map”, which corresponds to the planner in Figure 4b, stores all three-component sets of mobility and sensory state trajectories. Each trajectory of the set serves as the reference signal to one of the controllers in DIVA: the articulatory trajectory serves as input to the model-predictive controller, and the sensory trajectories serve as input to the respective auditory and somatosensory feedback controllers. The three-component representation of speech sounds in DIVA means that each speech unit has a fully-specified articulatory trajectory and time-locked sensory expectations. Uniquely among models discussed in the present paper, the sensory expectations are not generated online through an internal model, as in a state feedback controller.

The model predictive component of DIVA compares the predetermined desired position of the speech articulators at each point in time, **r**_*u*_(*t*), with their current predicted position, 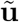, generating a control signal, 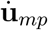. Implicitly, this assumes the existence of an internal model (not explicitly shown) that is able to predict the kinematic consequences of the motor commands with perfect accuracy. In order to generate the mobility state prediction, DIVA integrates the control signal over time. This enables comparison of the estimated state of the vocal tract articulators 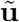 with the reference signal **r**_*u*_(*t*) independently of sensory feedback. Although the model-predictive controller is typically referred to as the “feedforward” controller in the DIVA literature, it is not a typical feedforward controller in the sense of “open loop” control traditionally described in control systems, because it relies on a comparison between the predicted current model state and a reference. In its current implementation, the predicted state also incorporates some auditory and somatosensory feedback information, as well, since those pathways converge with the model predictive pathway. However, if the auditory and somatosensory feedback controllers in DIVA are entirely removed, the model predictive controller would function appropriately in the absence of sensory information.

In the model predictive controller, the control signal is generated from the following equation: 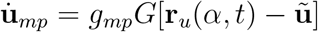, where *g*_*mp*_ is a scalar amplification gain applied to the motor command, and G is an additional gain that can be interpreted as a “go” signal, ranging between 0 (no movement) and 1 (maximal movement speed) as in Bullock and Grossberg (1988). Thus, the motor command is essentially a scaled version of an error signal, where the relevant error is between the articulatory reference signal and the predicted current position of the plant in mobility space. Note that the version of *u* that is used in computing the error signal is neither the true position of the articulators in mobility space, nor the one measured from the plant via sensory feedback, but an internal estimate of this state, 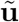. This estimate is generated by integrating the summed motor commands from all three controllers, and is equal to the motor position command issued to the plant. Effectively, the quantity 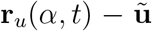 is an approximation of 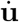 prior to scaling. The predicted current position of the plant is used purely as a way of converting the reference signal into a velocity, because the reference signal (a set of articulatory positions) cannot be used directly as a motor command (which must specify a change in those positions). Alternative ways of computing the motor command would eliminate the need for the model-predictive component of the feedforward controller, converting it into a true “open-loop” system. For example, the planner could approximate the first derivative of the entire articulatory plan, and issue that as the reference signal. Alternatively, the planner could issue the reference signal within a window surrounding the current time point, which would allow the controller to approximate the first derivative. Further details can be found in Appendix A.

The auditory and somatosensory feedback controllers closely follow the generic feedback control architecture. The auditory task space in DIVA is defined as the first three formants (F1-F3) and the somatosensory task space is defined as the positions of the individual articulators (proprioception) as well as the degree of contact between separate articulators (tactile sensation). Several publications have also envisioned the somatosensory space including representations of constriction locations and degrees, as in Task Dynamics (refer to sections describing Task Dynamics, below). The computations performed by the sensory feedback controllers in DIVA begin with a comparison between the reference signal and the sensory output of the plant to produce an error signal in sensory space. For the sake of simplicity, only the auditory feedback computations will be presented here, but the form is the same for the somatosensory pathways. The auditory error signal is defined as: **e**_*aud*_ = **r**_*aud*_(*α, t*) − **y**_*aud*_. This auditory task-space error is then transformed into a mobility-space error via the inverse kinematic equation: 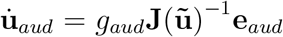. The matrix **J**(**u**) is known as the Jacobian, which provides a mapping between changes in mobility space and changes in task space. This mapping is dependent on the current mobility state (**u**) or, as in DIVA, a prediction of that state 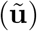. Specifically in DIVA, the Jacobian contains the rate of change for each of the dimensions of the task space for a corresponding change in mobility space. The matrix **J**(**u**)^−1^, is a pseudoinverse of the Jacobian, which allows for transforming task-space changes into mobility-space changes. The final motor command is then generated as the transformed error signal multiplied by a fixed gain, *g*_*aud*_. This represents a kind of proportional control, where the motor command, ignoring transformations for the moment, is simply a scaled version of the error signal. Further details can be found in Appendix A.

The output of the model predictive controller and sensory feedback controllers are summed to generate the final control signal, 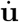. Thus, the final control signal passed to the plant is the velocity of the articulators (or 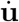) needed to produce the desired change in the position of the articulators (termed *motor movement command*). The control signal additionally includes the integration of 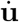 over time (**u**, or *motor position command*). This combined motor movement and position command is passed to the plant to drive changes in the position of the articulators. The plant also produces sensory outputs based on the position of articulators at each time point, **y**_*aud*_ and **y**_*somat*_. In DIVA, the output of the plant is in the space of the reference signal (F1-F3 for the auditory reference, position of the articulators as well as articulator contact for the somatosensory reference). This avoids needing to model an auditory or somatosensory perceptual system.

An important detail to note is that the auditory and somatosensory reference signals are specified not as unique trajectories with a single value at each time point, but as time-varying regions. The error signal for each space (auditory or somatosensory) is the distance from the current state to the edge of these regions. Thus, larger regions will allow greater variability in production, as no corrective error signal will be generated for any production that falls within the target region.

DIVA simulations have been able to qualitatively match human behavioral responses to auditory and mechanical perturbations (Guenther *et al.*, 2006; Tourville *et al.*, 2008; Villacorta *et al.*, 2007). The model has also been used to derive variable productions of /r/ (Nieto-Castanon *et al.*, 2005) based on a particular auditory target (low F3), a so-called “trading relationship” or “motor equivalence” where multiple articulatory configurations can be used for the same phoneme. Some older versions of DIVA that used time-invariant targets are also able to model carry-over and anticipatory coarticulation through the use of convex target regions (Guenther *et al.*, 1995).

Speech acquisition and learning have also received substantial consideration in the development of DIVA. The primary mechanism for learning within the model involves updating the motor plan based on generated auditory and somatosensory feedback motor commands. Details of this adaptive modification to the motor plan fall outside the scope of the present review. Nonetheless, this pathway is indicated by an open, labelled arrow in Figure 6.

**FIG. 6.**
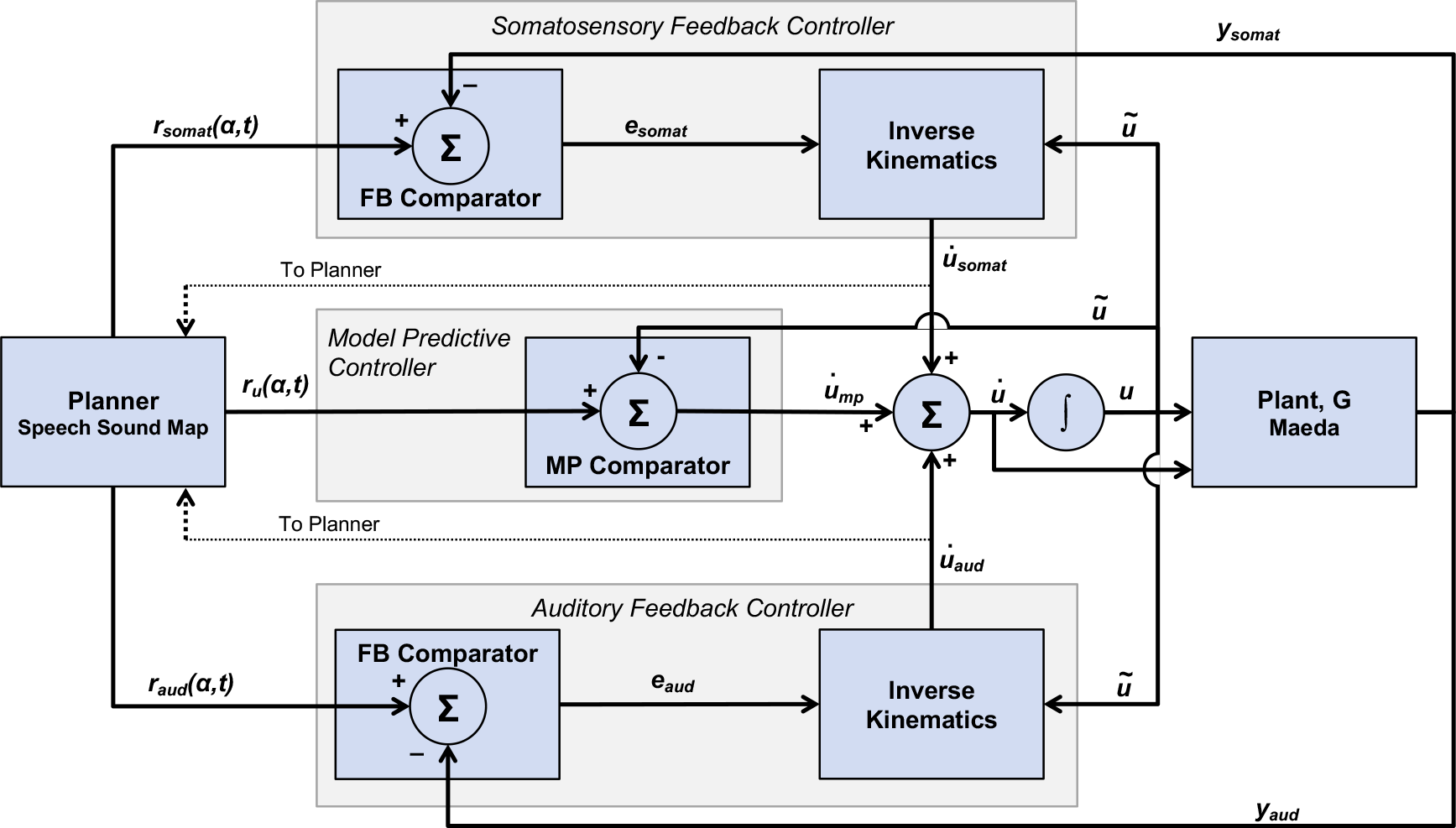
Control architecture of the DIVA model. The DIVA model has two feedback paths, auditory and somatosensory, that are schematically identical, and a model-predictive pathway. The feedback pathways compute an error between the planner’s signal and the output of the plant. This error is then used in conjunction with the state of the plant, *u*, to create a feedback control signal similar to the integrated model predictive-feedback control in Figure 5c. The model predictive pathway compares the desired position of the speech articulators with their current predicted position.

In addition to establishing the architecture of the speech motor control system, one of the primary motivations behind DIVA is establishing the neural basis of speech motor control. Individual components of DIVA have been mapped onto particular brain regions based on experimental neuroimaging results and model simulations (Bohland *et al.*, 2006; Ghosh *et al.*, 2008; Golfinopoulos *et al.*, 2011; Guenther *et al.*, 2006; Tourville *et al.*, 2008), and simulation studies have provided good matches to behavioral and neural activity recorded from human speakers during auditory and somatosensory perturbation experiments (Golfinopoulos *et al.*, 2011; Niziolek *et al.*, 2013; Tourville *et al.*, 2008; Villacorta *et al.*, 2007).

### B. Task Dynamics

The primary focus of the Task Dynamics model has been to model how invariant linguistic targets can generate continuous and context-dependent articulatory movements. The central hypothesis of this model is that articulatory movements are directed by the evolution of a task-level dynamical system whose invariant parameters are determined by the linguistic content of an utterance. TD was formulated by Saltzman and Kelso (1987) in general motor terms, and then by Saltzman and Munhall (1989) in the particular context of speech production (see Figure 7). TD is essentially a feedback control architecture, as described in Figure 7. The controller uses a feedback comparator to relate the desired state issued by the planner (**r**_*x*_(*α*, *t*)) to the current state of the system (**x**). On the basis of this comparison (**e**_**x**_), the controller computes a desired acceleration in task space 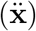 which is then transformed into a desired acceleration in mobility space 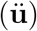. A crucial aspect of Task Dynamics is that both the desired state issued by the planner and the comparison performed by the controller occur in task space, not mobility space. This necessitates a transformation of the desired acceleration in task space into mobility space before it can be utilized as a motor command. The plant in the Task Dynamics model is the CASY model (Iskarous et al., 2003; Rubin et al., 1996), which is a geometric model of the vocal tract, similar in spirit to Maeda’s model.

**FIG. 7.**
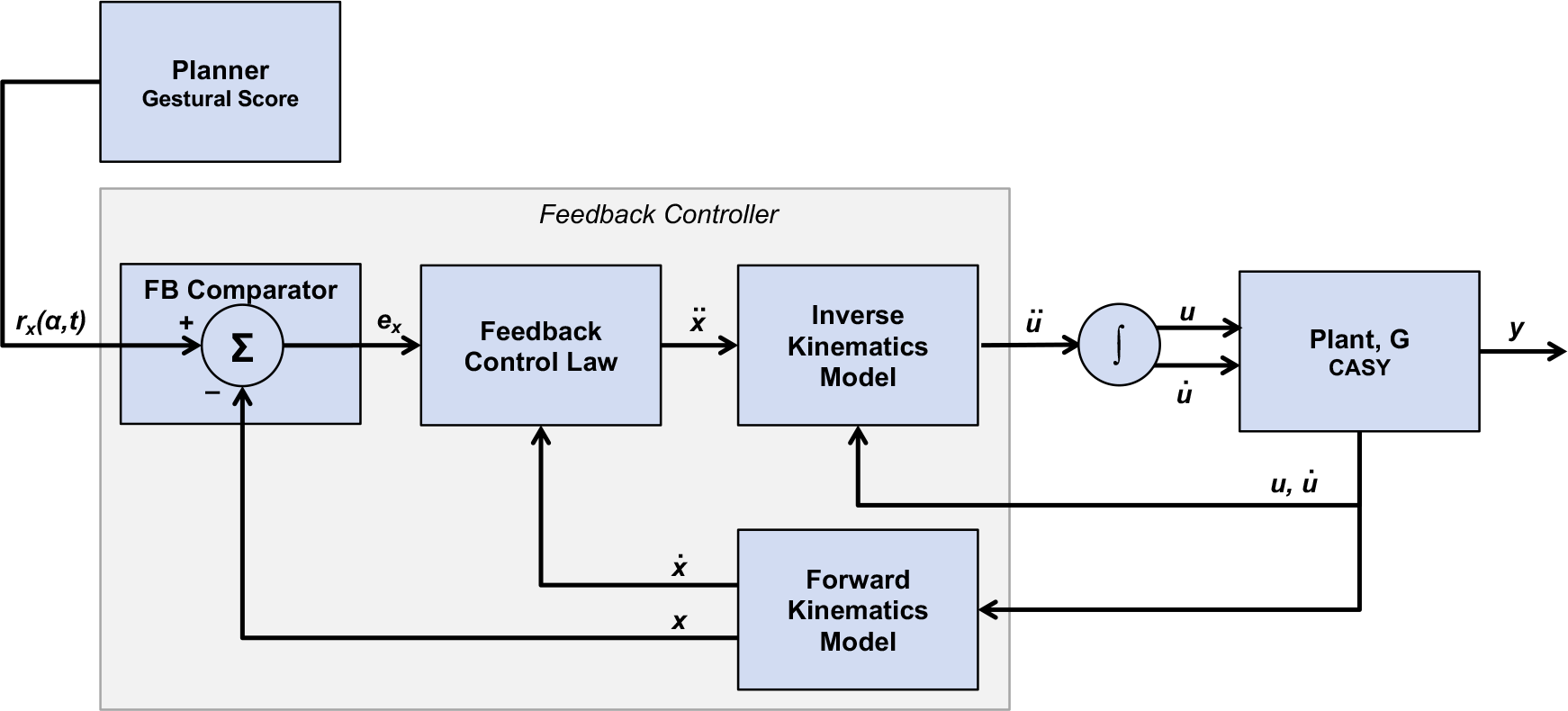
Control architecture of the Task Dynamics model. The system state, **x**, is broken out as both the state and change in state (first derivative), 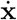. This information is used by the controller in the rectangle. Comparing this diagram to Figure 4b, one can see TD is a feedback control architecture.

One view represented in the literature and in the community of Task Dynamics is that it does not incorporate a feedback process. This misconception was perhaps most recently mentioned in print by Kröger and Birkholz (2007) who stated that a serious problem with the Task Dynamics approach has been the fact that “perception [presumably feedback] as a control instance for production is not considered”. Based on the discussion above, it should be clear that Task Dynamics is, in fact, primarily a feedback-driven system. One criticism that could be made of task dynamics, however, is that the model, as implemented, treats the feedback process as noiseless and instantaneous, which is overly simplistic. Given that the focus in task dynamics was on the development of the dynamic control law, this simplification would seem to stem from the specific emphases and interests of the authors, rather than some central conceptualization of speech motor control. Such was suggested by the authors in at least one publication (Saltzman and Kelso, 1987). It is also true that TD does not incorporate auditory feedback, which may, indeed, be a central property of the model. Similarly, the model assumes that the current state of the plant in mobility space is directly reflected via somatosensory feedback. Note that this is essentially the same assumption that DIVA makes, where part of the sensory feedback signal is simply the positions of the articulators.

The computations performed by the controller in TD begin with a comparison between the (task-space) reference signal and the task-space position of the plant to produce an error signal: **e**_*x*_ = **r**_*x*_(*α, t*) − **x**. The error signal is then used, along with the task-space velocity of the plant, 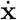, to update the task-space acceleration of the plant via the feedback control law (called the “forward dynamics equation” in the literature): 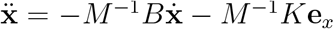, where M is a diagonal matrix of inertial parameters, B is is a diagonal matrix of damping coefficients, and K is a matrix of stiffness coefficients. Thus, the feedback control law takes the form of a second-order dynamical system that transforms the error signal into the second derivative of the task-space variable **x**. Since the task-space acceleration cannot be used directly as a motor command, it is necessary to transform this task-space acceleration into a mobility-space acceleration 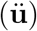. This is accomplished through the use of a pseudo-inverse Jacobian function: 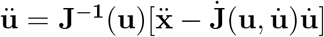. This mobility space acceleration can then be integrated to produce mobility-space velocity and position signals, 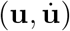, that can be used by the plant to drive changes in the position of the speech articulators. Further details can be found in Appendix A.

TD views speech motor control as a problem of point attractor dynamics. That is, motor tasks are conceptualized as points in task space, toward which the system is drawn by means of some governing control law which is a function of the system state. Task Dynamics describes the control law as a damped oscillator system (i.e., second-order dynamical system). Damped oscillator dynamics have a number of desirable properties in terms of defining a control law. In addition to the fact that damped oscillator dynamics are well-understood and easily characterized, the use of such dynamics to model task-directed behavior has the advantages that action patterns will be globally smooth and continuous.

TD is closely related to proportional-derivative control. It is common practice in engineering control systems to take integral or derivative information of the error signal into account (e.g., the ubiquitous proportional-derivative, PD, and proportional-integral-derivative, PID, controllers – e.g., Åström and Hägglund (1995)). Integrating the feedback error, for instance, allows a controller to recognize accumulated errors, which it can then attempt to nullify. Using the derivative of the feedback error, on the other hand, can minimize undesirable future trends in the error signal, such as overshoot, oscillation and instability. In PD control, the control signal **u**_*PD*_ is simply a weighted combination (given some weight matrices *K*_*P*_ and *K*_*D*_) of the error signal and its first derivative with respect to time: 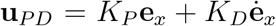. This equation looks remarkably similar to the feedback control law from TD: 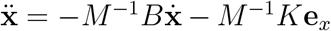, except that weights are specified, and 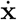 is substituted for 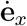. It can be easily shown that 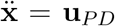, given that *K*_*P*_ = *M*^−1^*K* and *K*_*D*_ = *M*^−1^*B*, and knowing that **r**_*x*_ has a constant value, and therefore 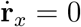. Thus, TD is equivalent to PD control up to the generation of the task variable acceleration signal, but differs in the additional transformation of the task space variables into mobility space, and integration of the mobility space variables.

The task space in TD is defined in terms of high-level articulatory tasks (in contrast to the positions of the individual articulators themselves). For speech, the tasks are suggested to be constriction actions (i.e., gestures) of the vocal tract, such as achieving closure of the lips, rather than the positions of the individual speech articulators (for the lip closure task, these would include the upper and lower lips as well as the jaw). A point attractor task is derived by the planner from a time-varying “gestural score” that issues the desired task state as a function of the currently active articulatory gestures. This definition allows TD to be easily put together with Articulatory Phonology (Browman and Goldstein, 1986). These two components form the basis for the perspective on speech production widely associated with Haskins Laboratories. Nevertheless, Task Dynamics and Articulatory Phonology are separate models that address different questions. Articulatory Phonology – proposed roughly in parallel with Task Dynamics – asserts that articulatory gestures are the primitive units of spoken language. Gestures themselves are conceptualized with AP as discrete vocal tract constriction actions, which can be composed into gestural “scores” that function as a motor program for a given utterance. Therefore, in broad terms, Articulatory Phonology addresses the question of how speech tasks should be defined, and how they can be composed into a motor program, whereas Task Dynamics addresses the question of how those tasks can be achieved and how that motor program can be realized in a physical system.

Use of second-order dynamics directly connects TD to research on action planning and execution in biological systems. For instance, the VITE model is an influential neural-inspired network model for explaining kinematic trajectory formation of directed movement (Bullock and Grossberg, 1988). VITE comprises a network of three interacting hypothesized neural populations, each coding a distinct quantity that is needed in the generation of the motor command, given some target position. These neural populations encode quantities related to the present position of the system, the desired target position, and the difference between the target and the present position. These interacting populations are configured in such a way that there are many structural similarities to the control architecture of TD. The result of these similarities is that the present position of a population will move in a way that is consistent with a 2^*nd*^-order dynamical system, much like Task Dynamics (as pointed out by, e.g., (Beamish *et al.*, 2006)).

One of the strengths of the model is accounting for coarticulatory effects. Coarticulation in this model is seen as arising from temporal overlap of independent and invariant articulatory gestures – the so-called coproduction model of coarticulation (Browman *et al.*, 1992, 1995; Fowler *et al.*, 1993). Other coarticulatory effects, such as clear vs. dark /l/ alterations, have been modeled at the planning level as changes in the temporal organization of gestures (Browman *et al.*, 1992, 1995; Zsiga *et al.*, 1994).

Very early results from the Task Dynamics model showed that it was capable of reproducing the compensatory behavior seen in mechanical perturbation experiments, where a lowered jaw position during production of a bilabial stop is compensated for by a higher lower lip and lower upper lip (Saltzman *et al.*, 1986). However, the model is unable to account for auditory perturbations, as there is no auditory feedback channel.

Task Dynamics can produce simple speech-rate effects by changing the dynamical parameters of the control law – e.g., by making the task-space motions more or less damped. In addition to these linear rate effects, the Task Dynamics model is able to produce a wide range of non-linear temporal effects seen in speech. Through the *π*-gesture model (Byrd *et al.*, 2003), the model is able to capture the non-linear slowing found adjacent to prosodic boundaries as well as capture many of the spatial effects, such as larger movements (Fougeron *et al.*, 1997), seen at those boundaries within a single framework. More recent work has extended the model to account for syllable structure and prosodic prominence (Saltzman *et al.*, 2008). While some recent work has started to explore neural mechanisms for some of the components of the model (Tilsen *et al.*, 2016), and a connection to the VITE neural model (Lammert *et al.*, 2018) has been established, the components of TD have not been explicitly related to specific neural structures.

### C. State Feedback Control

The State Feedback Control for speech production (SFC) model is a speech-specific instantiation of the general Kalman filter-type architecture in Figure 5c (Houde and Chang, 2015; Houde and Nagarajan, 2011). The primary focus of SFC has been to apply the insights gained from state feedback approaches in other motor domains to speech. This type of model is used widely in current theories of motor control in non-speech domains (e.g., work from Diedrichsen *et al.* (2010); Scott (2004); Shadmehr and Krakauer (2008); Todorov (2004); Todorov and Jordan (2002)), and is an evolution of a traditional feedback control system (Fig 4b). Recall that a primary challenge of feedback control is that sensory feedback is typically noisy and delayed, making the instantaneous state of the plant impossible to know with perfect accuracy. By adopting a Kalman filter-type architecture (Fig 5c), SFC presents, in a speech motor control context, one method by which sensory feedback may be integrated with internal model predictions to produce improved estimates of the state of the plant.

In the SFC model (shown in Figure 8), estimation of the plant state is done by an *observer* (refer to Fig 5c). This observer receives a copy of the outgoing motor command issued by the control law (also known as the efference copy) ^4^. Based on this signal, the observer predicts how the plant will move at the next time step 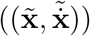 as well as the auditory and somatosensory feedback that will be received based on that predicted movement 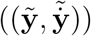. The predicted sensory feedback is then compared with actual sensory feedback to calculate a sensory error 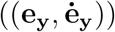. This error is then converted to a task state error (or task gain), via a gain function. Finally, the task state 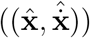 is estimated using the predicted state as well as the weighted sensory errors for both auditory and somatosensory predictions. As the gains associated with the sensory errors are assigned to optimize the final estimation, the observer in SFC functions is a Kalman filter (Todorov and Jordan, 2002), which provides the optimal *a posteriori* estimate of the state, under the assumption of linear processes of prediction and sensory feedback. Note that the sensory feedback the observer receives at any time point reflects the past state of the plant, while the state prediction reflects the current state. This delay is accounted for by delaying the sensory prediction before computation of sensory errors.

**FIG. 8.**
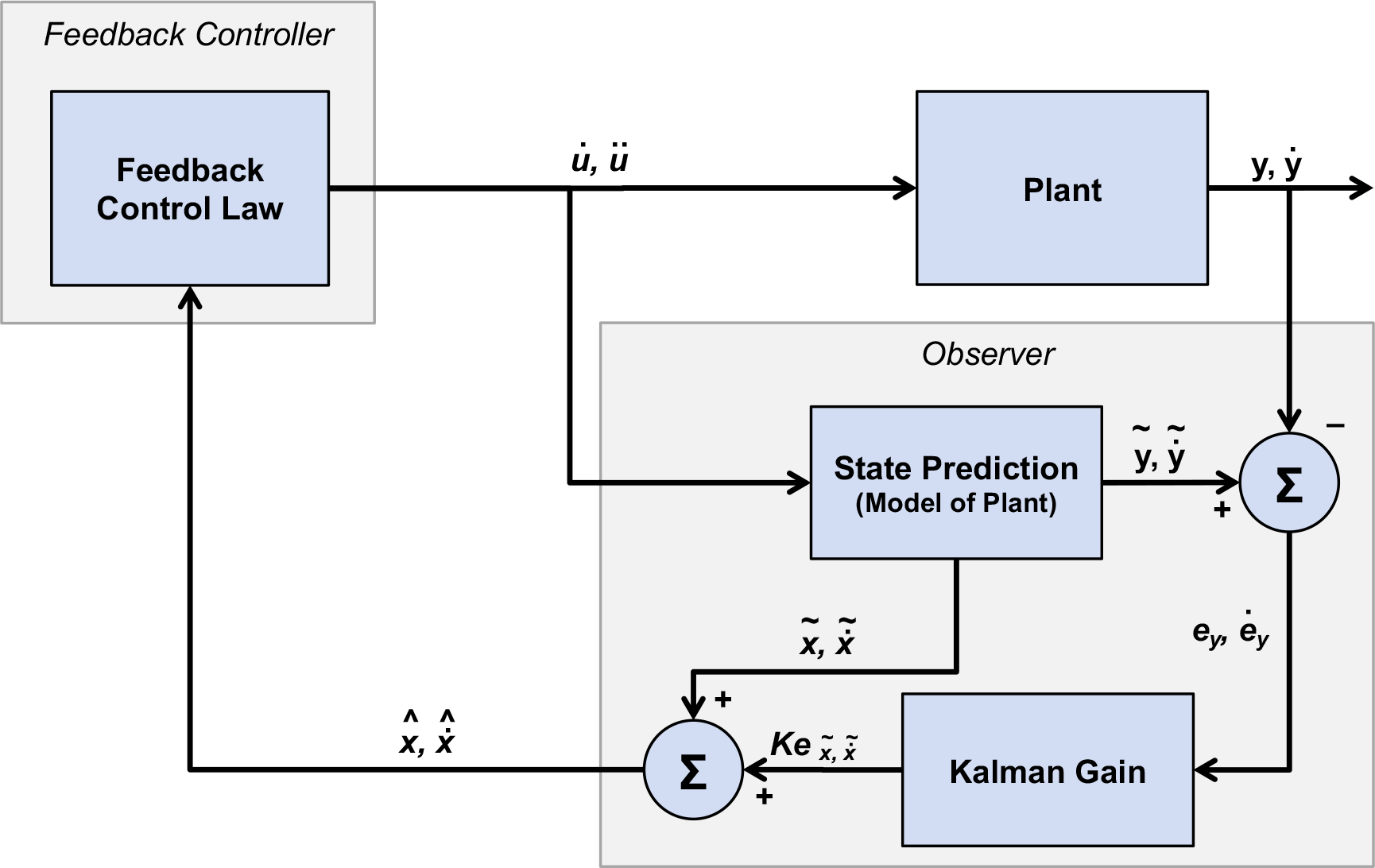
Control architecture of the State Feedback Control (SFC) model. The final state estimate passed back to the controller as a feedback signal, 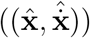, is derived from a combination of a state prediction process and sensory processes. Comparing this diagram to Figure 5c, one can see that SFC is an integrated model predictive feedback control architecture.

The model does not make explicit mention of a reference signal or a planner, and by extension does not make explicit mention of any comparison between sensory feedback and a reference. Providing a detailed description of the controller has not been a focus in the development of SFC, and therefore the controller, as presented in the literature, is represented by a generalized feedback control law which is a function 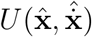 of only the state estimate. This control law could take almost any form. However, the authors of this review expect any feedback control law that produces reasonable speech production behavior would need to be a function of some kind of reference, whether an explicitly planned trajectory or a gestural score. Indeed, specifying the details of this feedback control law in SFC, and the addition of a planner module, have been a primary motivation for the development of the FACTS model, described below.

By combining a state prediction with sensory feedback to estimate the current state, the SFC model is able to act quickly by operating principally on an internal prediction of the plant state. This also allows the system to operate in the absence of sensory feedback, either when that feedback is too delayed to be of use (as for very fast speech movements) or when sensory feedback is unavailable (as when speaking in loud noise or in cases of non-congenital deafness). Yet, the system is still able to respond when the internal predictions do not match the incoming sensory feedback (either due to errors in the prediction process or due to external perturbations of the plant). Thus, this system combines the major advantage of traditional feedback control systems (robustness to perturbations) with that of feedforward control (fast, accurate movement even in the absence of sensory feedback).

Note that, in SFC as currently implemented, there is no distinction between task space and mobility space; they are effectively collapsed into a single space, such that commands are issued in task space. This means that the current implementation of SFC is only able to model a system where the goals of speech production are the same as the mobility space of the system. SFC has been implemented to control pitch, where the fundamental frequency of vocal fold vibration maps onto a one-dimensional mass-spring system.

This model has been shown to accurately reproduce the behavior patterns of human participants in pitch-alteration studies (Houde *et al.*, 2006). The model has also been shown to reproduce two neural effects seen in human speech: 1) the reduction seen in cortical electroencephalography (EEG) or magnetoencephalography (MEG) signals when speaking compared to listening to the one’s own speech played back over headphones or speakers (speech induced suppression) and 2) the enhancement of the EEG /MEG signals when seen when one’s speech is externally perturbed compared to when it is unperturbed (speech perturbation).

### D. FACTS

Recently, a new model – the Feedback Aware Control of Tasks in Speech (FACTS) model has been proposed that combines aspects of both Task Dynamics and State Feedback Control (Parrell *et al.*, 2006). Building on TD and SFC, FACTS combines elements of feedback control and model predictive control. FACTS is an attempt to combine the strengths of each model, while addressing the major shortcomings of each. Specifically, the Task Dynamics model includes a well-developed control law that relates the movements of the speech articulators to high-level tasks, but assumes perfect knowledge of the state of the vocal tract. Conversely, State Feedback Control focuses principally on how the state of the plant can be estimated from sensory information given the noise and time delays inherent in auditory and somatosensory perception, but has to date only been used to control a very simplistic one-dimensional model of pitch. FACTS combines the concept of state prediction and estimation from SFC with the planning model and vocal tract control of TD.

The architecture of FACTS is shown in Figure 9. The control component of the model is the same as that for the Task Dynamic model, with a planner generating a gestural score, which is passed to a feedback controller to generate changes at the task 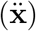 and mobility 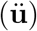 levels. This final motor command, 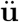, is passed to the plant to produce articulator movements as in Task Dynamics. However, where Task Dynamics passes the current plant and tasks states directly back to the feedback controller, FACTS uses an observer to estimate the task and plant states, as in the earlier SFC model. The final motor command 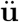 is passed to an internal model of the plant to generate predicted articulator positions 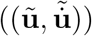, as well as auditory and somatosensory feedback 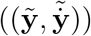. The estimated sensory feedback is then compared with sensory feedback from the plant to generate a sensory error 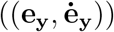. The estimated mobility state is generated from the predicted mobility state and the sensory error via an unscented Kalman filter, an extension of the linear Kalman filter to nonlinear systems (Wan and Van Der Merwe, 2001). The estimated mobility state is then converted to an estimated task state, needed by the feedback controller to generate the motor command at the next time point, via the same forward kinematics function as in Task Dynamics.

**FIG. 9.**
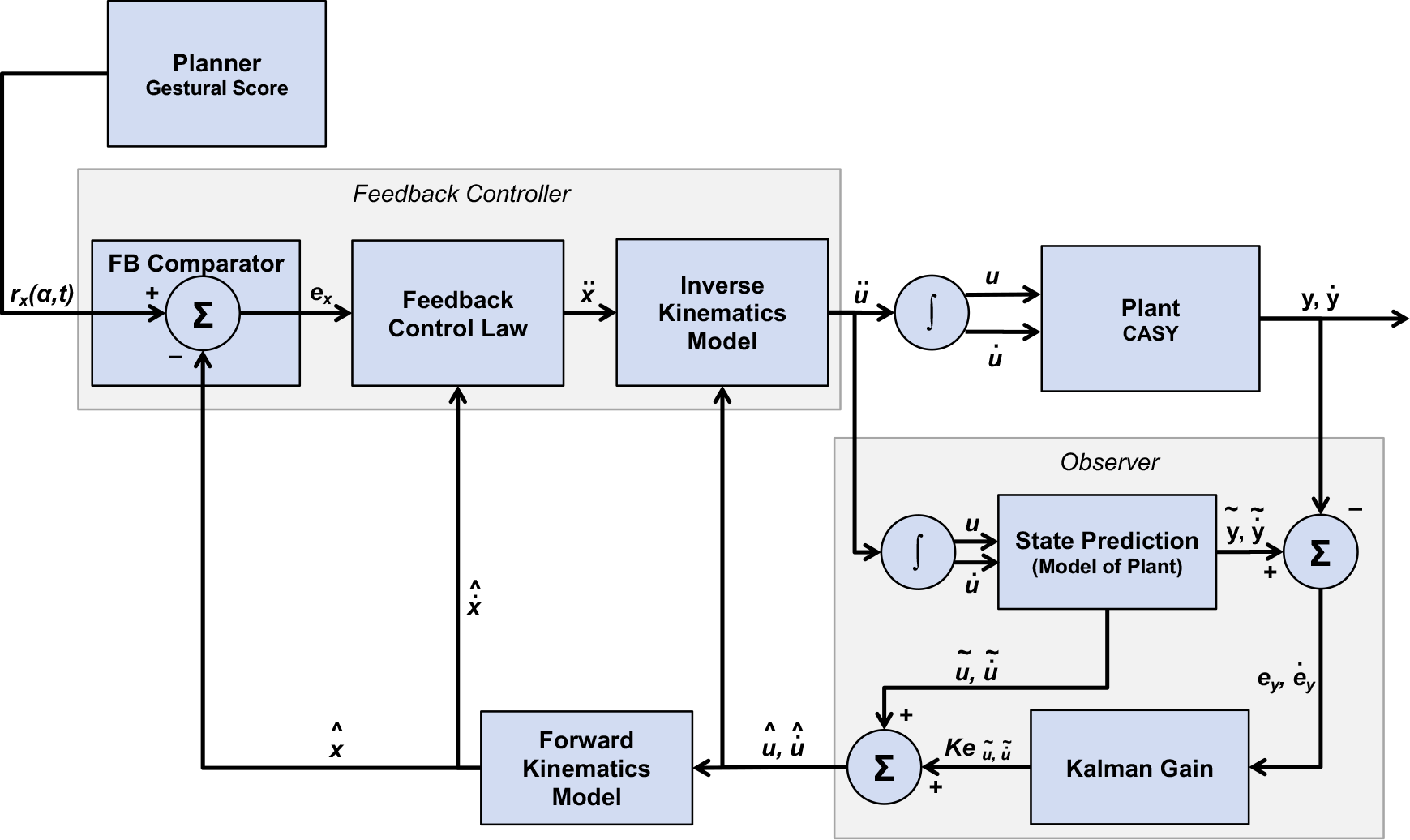
Control architecture of the Feedback Aware Control of Tasks in Speech (FACTS) model. FACTS builds upon the architecture of the Task Dynamics model by substituting an estimate of the mobility-space state for the true state through an observer module. The observer generates this mobility state estimate through a combination of an internal mobility state prediction and multisensory feedback. As such, FACTS is an implemenation of an integrated model predictive controller, like SFC.

The FACTS model is relatively new, and so remains mostly untested. However, the model is able to qualitatively reproduce human responses to external perturbations, including full compensation for mechanical perturbations and partial compensation for auditory perturbations (Parrell *et al.*, 2006). This partial compensation is a function of both auditory and somatosensory acuity. One of the features of FACTS is that it builds on the successes of the Task Dynamics model. Since many of the mechanisms of the controller are shared between the two models, FACTS can reproduce the successes of the Task Dynamics model, including coarticulatory effects.

### E. ACT

The primary focus in the the ACTion-based model of speech production, speech perception, and speech acquisition (ACT) model is the acquisition and development of speech motor control. Kröger *et al.* (2009) introduced ACT as a neurocomputational model that draws elements from both DIVA and Task Dynamics. The architecture of ACT, shown in Figure 10, is essentially a feedforward controller when viewed between the motor plan and the plant. DIVA-style dual auditory/somatosensory feedback pathways are also part of the model. However, those pathways feed indirectly to the planner, by way of high-level comparisons against abstract phoneme templates. Within the present framework, information used to modify the motor plan is considered to be part of the planner, and is therefore outside the scope of low-level control, as defined here. This pathway is indicated by an open, labelled arrow in Figure 10. The plant in ACT is a three-dimensional kinematic model with articulatory control parameters similar to the Maeda and CASY models (Birkholz *et al.*, 2006).

**FIG. 10.**
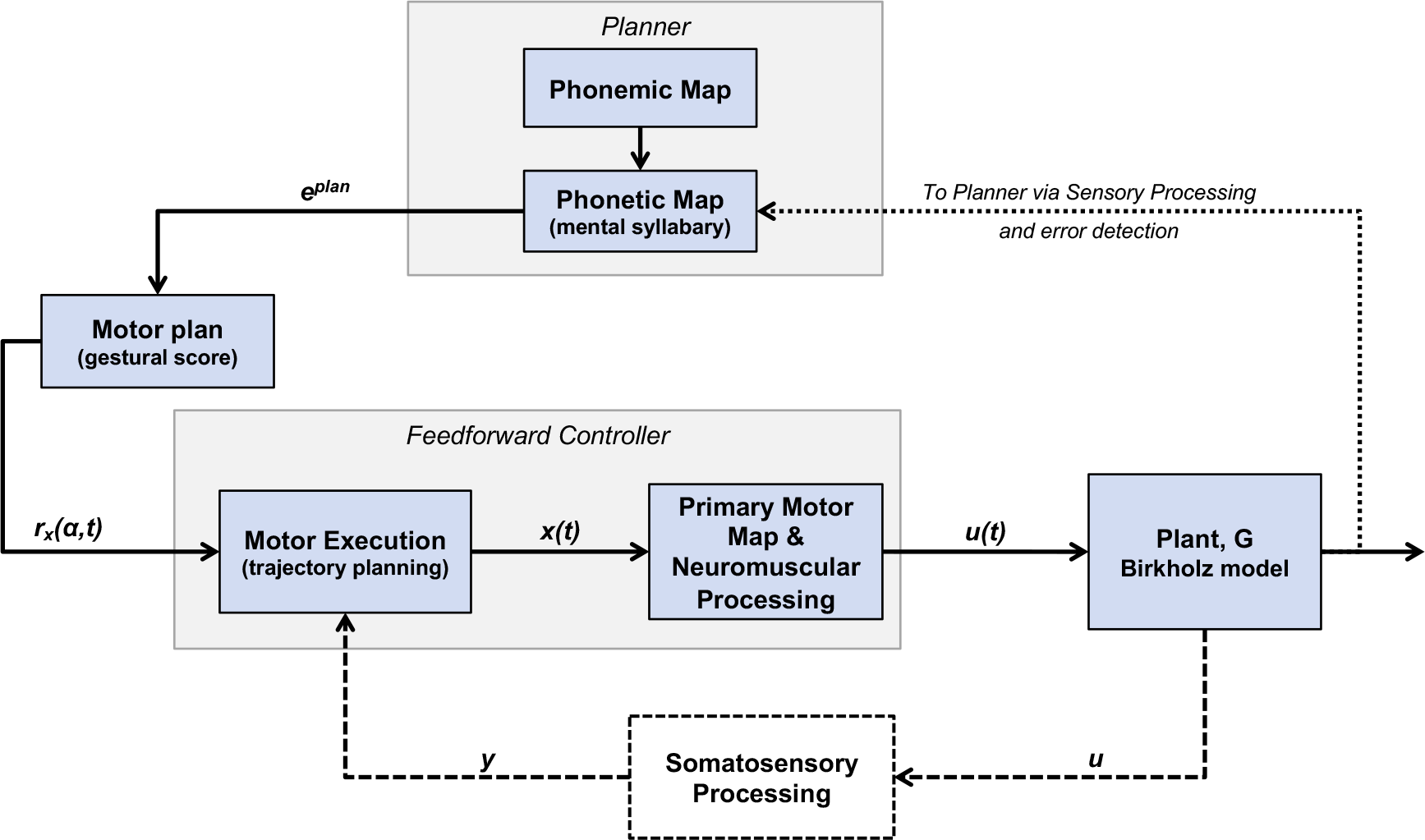
Control architecture of the ACTion-based model of speech production, speech perception, and speech acquisition (ACT model). ACT draws from both DIVA and Task Dynamics for its architecture, with the model comprising both feedforward and feedback pathways (both somatosensory and auditory), but relying on point-attractor dynamics for its reference signal.

The planner in the ACT model relates to both the speech sound map of DIVA and the gestural score in the Task Dynamics model. Like in DIVA, the basic unit of speech is assumed to be the syllable, and each syllable is represented by a model neuron in the phonemic map (cf. the speech sound map in DIVA). As in DIVA, these abstract syllable representations are linked to specific motor and sensory plans. This is accomplished in ACT through the phonetic map. Unlike in DIVA, where the motor plan is represented as a time-varying desired articulatory position signal, the motor plans in ACT are defined in terms of high-level dynamic tasks (or gestures) as in Task Dynamics. Each motor plan is, in effect, a gestural score, which defines the activation levels and temporal extent of each speech gesture, with each speech gesture being defined as a dynamical point-attractor (**r**_*x*_).

The phonetic map, in addition to linking the syllable to the motor plan, also links the syllable to associated sensory (auditory and somatosensory) expectations. One conceptual difference between ACT and DIVA is that DIVA views the sensory plans as the targets of speech that have associated motor plans, while in ACT the targets are the high-level task gestures with associated sensory expectations. This conceptual difference is reflected principally in terms of how the models are trained (an issue not taken up within the scope of the present review), but the basic architecture of the models is essentially the same: a high-level syllable activates a motor plan used for feedforward or model-predictive control and a sensory plan which can be compared against afferent sensory information.

The core control architecture in the ACT model borrows ideas from Task Dynamics, but is quite distinct. As discussed above, TD makes use of task-space comparisons between a reference, derived from the task-based gestural score, and the current (somatosensory) system state to control task-space movements given a control law that is consistent with damped oscillator dynamics. ACT, on the other hand, uses the reference, similarly derived, to directly drive motor action in a feedforward fashion. This is accomplished by the motor execution module, which uses the reference **r**_*x*_(*α, t*) to generate a trajectory in task space (**x**(*t*)) that is consistent with damped oscillator dynamics. The task-space trajectory must be transformed into a mobility-space trajectory (**u**(*t*)) that can be used as a control signal to drive movements of the plant. This transformation is accomplished by the primary motor map. A subsequent neuromuscular processing step exists in the model, and is presently implemented as a direct, linear mapping. Plans exist for this component to eventually map control signals onto individual and/or combined muscle groups in a neuromuscular model. An additional pathway for somatosensory feedback processing is also planned. This is indicated by dashed lines in Figure 10. This feedback pathway, included in published figures representing ACT, would be used to “control motor execution”, presumably in a fashion similar to DIVA. This pathway has not yet been implemented, and the details of its properties have not been fully developed.

Like DIVA, the ACT model also has dual somatosensory and auditory feedback pathways. The principal way these feedback pathways are used in the model is to compare the current state of the plant against pre-learned templates representing the desired somatosensory and auditory states. A crucial difference between ACT and other models is that this error signal is used to influence the motor plan, rather than as part of the controller. That is, sensory feedback is used to detect sensory errors for updating the phonetic map to drive trial-to-trial adaption, a model of development and learning.

One difference between the ACT model and others is that the mappings that relate the different signals (syllables↦**r**_*x*_, syllables↦**r**_*y*_, **r**_*x*_↦**r**_*u*_, 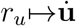, etc.) are implemented via tunable neural networks rather than as closed-form mathematical expressions. These networks are tuned during a learning phase. Some versions of DIVA presented in the literature, especially earlier in DIVA’s development, had neural networks involved in these mechanisms (Guenther, 1994). The use of trained neural network models for these transformations allows for flexibility in the form of the transformations. It opens the possibility that the transformations might take forms that deviate in unexpected, and potentially even biologically-plausible, ways when compared to mathematically-driven transformations typically adopted. The use of neural networks also makes it likely, however, that key transformations, such as the control law and the inverse kinematic transformations, cannot be easily written down analytically in closed form.

The ACT model is able to produce motor equivalence in articulators linked to the same gesture due to the use of high-level tasks rather than articulatory positions as the basic unit of the motor plan (Kröger *et al.*, 2009). The model is also capable of adaptive learning based on high-level auditory errors or somatosensory perturbations, by changing the motor plan. However, the lack of feedback pathways in the controller means online compensation to these perturbations is not accounted for. The ACT model includes hypotheses about the neural structures that underlies the different components but to date has not been used to generate simulated neural activity to compare to neural data.

### F. GEPPETO

The GEPPETO (GEstures shaped by the Physics and by a PErceptually oriented Targets Optimization) model (Patri *et al.*, 2015; Perrier *et al.*, 1996, 2005;) is a model of speech control based on the equilibrium point hypothesis (Feldman, 1986). The primary focus of GEPPETO has been to investigate the hypotheses that 1) targets for speech production are discrete and phonemic, 2) biomechanics plays a non-trivial role in speech motor control, and 3) speech motor control employs optimal planning principles. In GEPPETO, as in the equilibrium point hypothesis, control occurs at the level of individual muscle lengths. Thus, the mobility space in GEPPETO is composed of lengths, *u*_*k*_, of individual muscles *k*. The command generated by the central controller is a muscle length, or threshold, above which motor neurons will be recruited to contract the muscle. This threshold length is known as the equilibrium point or λ. Afferent feedback from the muscle about the current muscle length is compared against the current λ, and contractile force is generated if the muscle length is above the threshold. In GEPPETO, the activation (*A*) of each muscle at time t is based on both the current muscle length *u* and the current change in muscle length 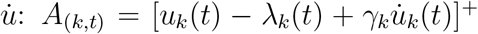, where γ is a damping parameter that stabilizes the system. Muscle activation is only generated when the muscle length is greater than λ: [*A*]^+^ = {*A*, *if A* >= 0; 0, *otherwise*}.

The muscle activation generated by the feedback controller then leads to the generation of force (*f*) in the individual muscles at the level of the plant: *f*_*k*_(λ_*k*_, *t*) = *ρ*_*k*_[*exp*(*c*_*k*_*A*_*k*_(λ_*k*_, *t*) − 1)],where *ρ* is a magnitude parameter related to the cross-sectional area of the muscle and *c* is a feedback gain. In this feedback control architecture, force can be generated either by changes in the current λ or by changes in the length of the muscles. Importantly in this approach, the ultimate position of the plant results from a combination of descending control (λ values), plant biomechanics, and physical constraints.

The GEPPETO model, shown in Figure 11, combines the low-level feedback control structure of an equilibrium point model with a high-level feedforward controller that takes acoustic speech targets, defined as convex regions in acoustic (F1-F2-F3) space, as input and output λ values that are passed to the feedback controller. Thus, the task space for GEPPETO is acoustic in nature (though see for a recent extension of the model to additionally include somatosensory targets). Critically, given the emphasis on the physics of the speech plant, GEPPETO uses a dynamical biomechanical model of the plant with control occurring at the level of muscles rather at the level of geometric model parameters/articulators as in the Maeda or CASY plant models. Most published papers on GEPPETO include only the tongue as a controllable articulator. It is modeled as a finite-element model with six muscles whose lengths can be independently controlled. The other vocal tract surfaces and articulators are fixed.

**FIG. 11.**
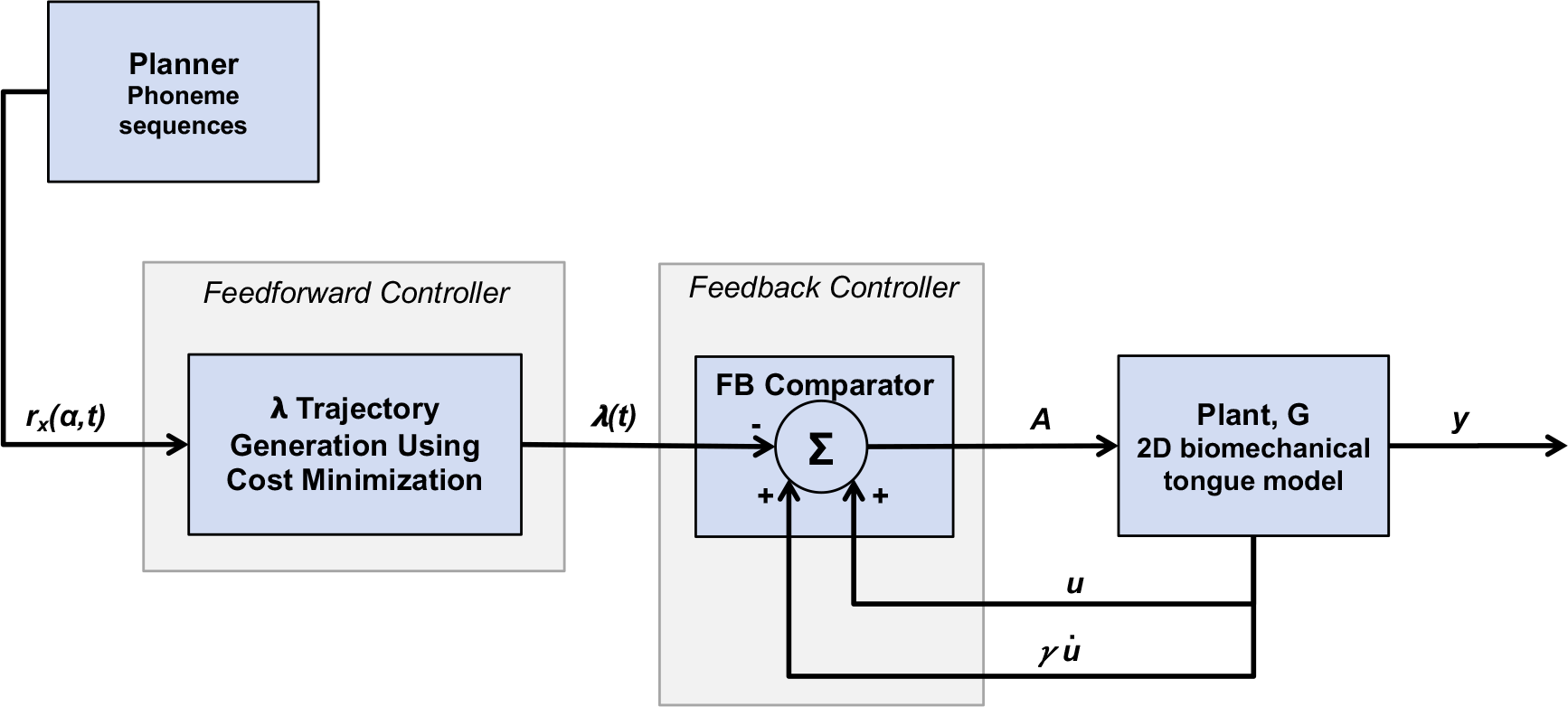
Control architecture of the GEPPETO model. GEPPETO is based on the equilibrium point hypothesis, employing feedback control at the level of individual muscles, with relatively realistic biomechanics to move the speech articulators. Control is mediated by a feedforward process that transforms acoustic speech targets into equilibrium point values.

The output of the planner in GEPPETO is a series of *n* acoustic speech targets (*ϕ*^1^, …, *ϕ*^*n*^), each of which has an intended duration (*T*^1^, …, *T*^*n*^). This duration can be affected by variables such as speech rate or stress. An additional constraint sets the amount of effort to be used for each speech target (*w*^1^, …, *w*^*n*^), where effort is based on the amount of force that will be generated to produce that target across all the muscles of the plant, categorized into three levels: *w* ∈ {“*weak*″, “*medium*″, “*strong*″}.

This time series of targets **r**_*x*_(*t*) = {(*ϕ*^1^, *w*^1^, *T*^1^), …, (*ϕ*^*n*^, *w*^*n*^, *T*^*n*^)} is then passed to the feedforward controller to generate a time series of λ values for each of the six muscles in the plant, (λ_1_(*t*), …, λ_6_(*t*)). These λ trajectories are generated for each utterance using an optimization procedure that minimizes displacements in mobility space (i.e. changes in muscle lengths) while producing tongue movements that will achieve the required acoustic targets at the required time with the required amount of effort. In this optimization process, learned internal models are used to estimate the amount of force and acoustic signal generated for any given motor command.

GEPPETO shares certain characteristics with other models. First, speech goals are defined as regions in acoustic space (F1-F2-F3), as in DIVA. Second, feedback signals are never directly compared against the output of the planner, as in ACT. GEPPETO differs in key ways from other models, however. First, the speech targets in GEPPETO are hypothesized to be discrete in time, rather than time-varying regions as in DIVA. Second, the feedforward and feedback controllers in GEPPETO are arranged in a unique, serial or hierarchical arrangement, such that the output of the feedforward controller is used as the input to the lower-level feedback controller. Third, unlike the preplanned trajectories in DIVA, GEPPETO generates new movement plans for each utterance. Finally, it is notable that GEPPETO is unique in the fact that the plant’s inputs are not given in mobility-space variables.

The largest success of the GEPPETO model has been to replicate many of the characteristic kinematic patterns of speech movements, including velocity profiles (Payan *et al.*, 1997), tongue loops in velar stops (Perrier *et al.*, 2003), and the relationship between velocity and movement curvature (Perrier *et al.*, 2008). This work shows that many of these phenomena need not be directly controlled, since in GEPPETO they are emergent properties of linear changes in λ values. One of the drawbacks of the optimization approach in GEPPETO is that it produces identical trajectories each time the same utterance is produced, unlike the variability seen in natural speech. Recently, however, the GEPPETO model was expanded by implementing it in a probabilistic Bayesian framework (B-GEPPETO) that is able to account for token-to-token variability (Patri *et al.*, 2015;). This newer model also incorporates somatosensory phonemic targets in addition to auditory targets.

### G. Other models

All the above models include, at a minimum, the ability to generate motor commands based on some motor plan. These motor commands are then used to move a vocal tract model of some kind. While such complete models are the primary focus of the current review, it is important to also mention more conceptual models which have not been implemented to the same degree. The Hierarchical State Feedback Control model (HSFC) (Hickok, 2012a,b, 2014) is an attempt to combine speech motor and psycholinguistic approaches to speech production. It is a version of an integrated predictive/feedback controller, sharing some aspects with the State Feedback Control model of speech production (Houde and Nagarajan, 2011). Tian & Poeppel (Tian and Poeppel, 2010) propose a hybrid model predictive/feedback control model of speech motor control. The overall architecture is also very similar to the State Feedback Control model (Houde and Nagarajan, 2011).

A few other models of speech motor control have been proposed that have focused primarily on the biomechanical properties of the plant rather than on the control architecture per se (Dang and Honda, 2002, 2004; Laboissiere *et al.*, 2018; Ostry *et al.*, 1996; Perrier *et al.*, 1996; Sanguineti *et al.*, 1990). While these models do not relate control to linguistic speech targets (i.e. describe how or why certain muscle contraction patterns would be used), the success of these models in recreating measured articulatory trajectories deserves mention in the context of the present review.

One class of these models (reviewed in Sanguineti *et al.* (1998), is based on the equilibrium point control. While this is the same general approach as taken by the GEPPETO model, the focus of this work differs. Rather than implementing control of the speech motor system in terms of higher-level linguistic or task-directed (auditory, articulatory) control, these models focus principally on how muscle forces are generated to move the speech articulators. Typically, the goal is to drive movements to match measured human speech kinematics. These models essentially implement a feedback controller, albeit one that functions entirely at the level of the plant without any distinction between task and mobility space. A separate set of biomechanical models assumes that muscle activations are the output of the controller, rather than equilibrium points (Dang and Honda, 2002, 2004). This is a purely feedforward control architecture.

Both the equilibrium point models (Sanguineti *et al.*, 1998) as well as the direct activation models (Dang and Honda, 2004) have been shown to fit articulatory data well using similar biomechanical models. Interestingly, results from both models suggest that motor commands to certain muscles (or muscle groups) will drive the speech articulators towards a similar location regardless of their initial position. This suggests that speech motor control may be simplified by the use of muscle synergies that will drive the system to a target spatial configuration without the need for complex inverse dynamics models that calculate the precise muscle activations needed for each individual movement.

One important thing to note is that, because they focus on the generation of muscle forces given some given motor commands, this class of models is generally complementary to and compatible with control models that output motor commands as articulatory positions, and ignore the generation of muscle activations (such as DIVA, TD, ACT, and FACTS). With some modifications, the output of these models could serve as the input to an equilibrium point model or the Dang & Honda model. In fact, equilibrium point control has been implemented within the DIVA architecture (Zandipour *et al.*, 2004).

## IV. DISCUSSION

The primary goal of the current paper has been to clearly lay out the architectures of a crucial component of existing speech motor control models: the control layer (see Figure 1), that attempts to produce accurate tracking of speech articulation kinematics given a motor plan. Common terminology and basic principles of motor control were used to describe each model, to understand the commonalities between these models, as well as how they differ. It was shown that these models can be cast as special cases of generalized feedforward (Fig 4a), feedback (Fig 4b), and model predictive (Fig 4c) controllers. The models discussed here differ in which of these components are used (e.g., some are lacking either feedforward or feedback elements of control), and in the detailed implementation of these mechanisms. These differences are summarized in Table I. Speech production is, however, a complex process with many additional and important considerations, including higher-level motor planning, linguistic, communicative and even social considerations, as well as learning and developmental aspects, all of which contribute to the wide variety of speaking styles observed in real human speech. These aspects are beyond the scope of the present paper, but would make an interesting subject future reviews.

**TABLE I.** Summary of which aspects of motor control modeling are present in each model.

|  | DIVA | TD | SFC | FACTS | ACT | GEPPETO |
| --- | --- | --- | --- | --- | --- | --- |
| Feedback Pathway | Y | Y* | Y | Y | Y | Y |
| Feedforward Pathway | N | N | N | N | Y | Y |
| Internal Prediction/State Estimation | Y | N | Y | Y | N | N |
| Principal Reference | Tourville and Guenther (2011) | Saltzman and Munhall (1989) | Houde and Nagarajan (2011) | Ramanarayanan *et al.* (2016) | Kröger *et al.* (2009) | Perrier *et al.* (2005) |

There are clear differences among models in terms of how their final execution of speech motor control is influenced by feedback signals originating from the plant. ACT, for instance, incorporates no explicit feedback into its control mechanisms. SFC implements proportional control, meaning that the motor commands are linearly proportional to the feedback error. DIVA’s also implements proportional control which, for its hybrid architecture means that motor commands are linearly proportional to both the error (in the feedback pathway) and the reference (in the feedforward pathway) signals. The simplicity of these designs relative to common engineering approaches is notable. As mentioned above, and by way of example, engineering control systems often take information about the integral or derivative of the error signal into account in order to provide quicker convergence to the target and to deal with persistent errors, respectively. TD – as well as FACTS, by way of adopting key control elements from TD – provides slightly more complexity through a form of PD control, albeit not strictly in the traditional engineering sense of PD control.

A related distinction between the models under consideration is how they function in the absence of feedback. TD, for instance, is solely a feedback architecture, and cannot function in the absence of feedback signals. Similarly, GEPPETO would not be able to function in the absence of proprioceptive feedback about muscle length. Other models could continue to function without feedback. DIVA is a hybrid feedback/model predictive architecture that could rely exclusively on its model predictive mechanisms to generate motor commands in the absence of explicit feedback. With the presence of feedback signals, SFC and FACTS can utilize that feedback to produce optimal or near-optimal state estimates (under certain strong assumptions, such as linearity of the plant (Kalman *et al.*, 1960)), but in the absence of feedback can still rely on the internal state prediction component of their broader state estimation process to continue functioning through model predictive control. ACT is a purely feedforward architecture that can function as designed in the absence of sensory feedback. However, this also means that it is not sensitive to sensory feedback, unlike the human speech motor control system.

Among models that incorporate feedback, one of the most basic differences is whether certain feedback signals are treated as idealized signals that are directly and instantaneously observable, or whether they are treated as true sensory signals that may be potentially noisy/delayed, subject to conditioning by internal models and that correspond with known neurological signals. While it seems intuitively correct that any model of biological motor control should focus on the latter, the former has been sometimes intentionally chosen in specific aspects of the models, in the interest of focusing on other aspects of control. TD provides only an idealized view of feedback concerning the positions and velocities of articulators that does not model the sensory processes in any meaningful way. DIVA, TD and FACTS also make simplifying assumptions about the somatosensory feedback signal, which is assumed to be more or less equivalent to the plant’s mobility variables. DIVA’s auditory and somatosensory feedback are slightly less idealized in that they correspond to known, independent neurological pathways and can incorporate delays associated with sensory transduction and processing. SFC and FACTS begin with the assumption that sensory feedback will be noisy and/or inaccurate, and use that assumption to motivate the well-elaborated integration of sensory feedback with internal model predictions to provide more accurate estimates of the state of the plant. GEPPETO provides perhaps the most realistic implementation of somatosensory feedback given that the feedback in the model (muscle length and change in muscle length) corresponds to well-known afferent signals from muscle spindles. However, no current models seriously attempt to model the sensory system itself – they take it as given that critical information (e.g., formants, articulatory positions) can be extracted from the raw sensory input.

Most models are purely kinematic in how they approach control, in that motor commands are stated in kinematic terms (i.e., as state configurations, and not as forces) and do not account for dynamical considerations related to the effects of inertia, centrifugal and centripetal forces, and the effects of gravity. Control systems that are strictly linear, rigid and slow-moving, highly damped, or that have specialized designs can sometimes operate purely kinematically. It seems likely, however, given existing literature (e.g., Derrick *et al.* (2015); Ostry *et al.* (1996)) that such considerations may be non-negligible for speech production in the biological case. A kinematic approach can be, in the opinion of the authors, partially attributed to models of the plant used in most speech motor control models, which are nearly all kinematic in nature. It is worth noting that other plant models are attempting to provide enhanced biomechanics (Derrick *et al.*, 2015; Gick *et al.*, 2011; Lloyd *et al.*, 2012) as well, even if a full review of biomechanical vocal tract models is beyond the scope of the present review. GEPPETO represents a notable effort to move beyond kinematic treatment of control, and of the plant, by incorporating a mobility space that represents muscle lengths, as well as motor commands that represent muscle activations that are used to generate muscle forces in a relatively realistic biomechanical model of the tongue.

All architectures rely on a motor plan of some kind – whether an explicitly planned trajectory or a gestural score – that is formed at a higher level of motor processing, and which is issued to the controller in order to be carried out. SFC is a partial exception to this general statement in that, as mentioned above, that model does not explicitly mention the incorporation of a plan, even though the generalized structure of its controller would be able to incorporate a planning module if more detailed specification required it (a specification which FACTS has subsequently elaborated upon). Models of speech motor planning have been discussed and elaborated upon in the literature (Bohland *et al.*, 2010; Byrd *et al.*, 2009; Civier *et al.*, 2013; Saltzman *et al.*, 2008), and display a surprising amount of variety. Although the planning level is beyond the scope of this paper, it is worth noting the variety of planning mechanisms that have been proposed in order to help narrow some of the longest-standing debates concerning speech motor control. In particular, drawing a clear distinction between control architectures and planning mechanisms, as this review has attempted to do, makes it apparent that much of the debate over the quality of competing models of speech production would appear to be concentrated at the planning level, and not at the level of control. For instance, issues surrounding the nature of production goals (e.g., acoustic vs. articulatory) and the composition of those goals into utterance-size units would primarily be a concern of the planning level. Any role for muscle synergies (Ramanarayanan *et al.*, 2014) and motor primitives would be most naturally incorporated into the planning level, and not the level discussed in this review. The nature of speech production goals has been the subject of particularly strong debate for decades, and is reflected in the nature of the feedback and reference signals in the models, which may be auditory and/or somatosensory, as in DIVA and SFC, or articulatory, as in Task Dynamics. Interestingly, the nature of the feedback signals would appear to have little bearing on the specific architectural choices of the models – the architectures being general enough to handle a range of signals without substantial changes to their configuration.

The parameters that determine the overall characteristics of control are time-invariant in most current models, thereby limiting the models in their ability to capture specific aspects of behavior that require those parameters to change over time. Models may struggle, for instance, to account for interspeaker differences, or long-term changes in speech motor control that occur during development and aging, that could be modeled by adjustments to control parameters. Controllers that adapt their parameters over time are the subject of adaptive control (Åström and Wittenmark, 2013). This well-studied branch of control theory may provide a foundation for models of speech production to incorporate such parameter adjustments as a way to represent the mechanisms of differences or changes mentioned above. A full treatment of adaptive control is outside the defined scope of the present paper, as are issues surrounding speech development. Nonetheless, it should be noted that inroads into adaptive control have been made by some of the models discussed here. ACT allows for motor planning to be adapted based on sensory feedback errors. DIVA, too, adapts planned trajectories based on the feedback controller output. This adaptation is of primary importance during development, but can lead to changes at any time.

Shorter time-scale cognitive and physiological factors – for instance, due to attention, fatigue and motivation – as well as stochastic variability (Munhall *et al.*, 1994; Saltzman *et al.*, 1995; Tilsen, 2017) may also most naturally be handled through adjustments to control-level parameters. Efforts have been made to model learning and adaptation at the planning level (e.g., GODIVA). However, the value of the proportional gain in DIVA’s controller, as well as the weights assigned to the feedback and model predictive pathways in their contribution to the motor command, are assumed to be fixed in fully adult speech. Similarly, the damping and stiffness parameters of the controller in TD are fixed in value. A notable counterexample to this generalization comes from Kalman filter-based architectures, such as SFC and FACTS, which change the weight assigned to sensory feedback and internal model predictions, toward combining them into a single state estimate, based on the degree of statistical reliability of those two pathways. Such adaptation may be useful in modeling the impact of sensory feedback impairment on speech motor control. Another notable example of this type is DIVA’s GO signal, which can be adjusted by higher-level processes in order to control the initiation of movement and overall speaking rate.

A clear understanding of how the various models are structured can aid in clearly defining theoretical questions of interest. For instance, the many similarities of the models discussed in this review naturally raise questions about what is gained by allowing the remaining model dissimilarities to persist, and whether the models can converge to a single, unified model of the control layer in speech motor control. There is no mathematical reason why the feedforward/feedback pathways embodied by DIVA couldn’t be combined with the forward dynamic control of TD, as well as the feedback/internal model-based state estimation in SFC. Indeed, FACTS, as a combination of complementary elements of TD and SFC, has already taken a step toward beginning these potentially useful combinations. Whether such a unification is sensible from a theoretical point of view, and precisely what form that unification might take, can be stated very precisely in mathematical terms using the model architectures. In general, models can help in defining and circumscribing the space of possible architectures and solutions to a specified biological control problem (Schaal and Schweighofer, 2005).

A related, empirical question is whether a model unification is useful for explaining observations from human speech data. Among the many benefits of developing formal models of speech motor control is that models can be used to make specific, quantitative predictions about human speech behavior that are testable in light of data. The predictive capabilities of formal models can also guide the design of new experiments to test specific aspects of theory and modeling, inspired by the behavioral predictions of the model, and perhaps piloted *in silico*. Empirical questions regarding the models need not be limited to observable behaviors, either. Models can also facilitate clearer connections to be drawn between specific model mechanisms and their observed neurological counterparts, either through structural or functional neuro-imaging. The connection between engineering and biological mechanisms has been well developed in several domains of motor control, including speech motor control (Guenther *et al.*, 1998) and oculomotor control (Lisberger, 1988; Robinson, 1981; Shibata and Schaal, 2001).

The utility of speech motor control models additionally extends beyonds clarifying and formalizing our understanding of speech motor control itself. Models can also be useful for practical applications in speech synthesis. Control models, coupled with faithful mechanical models of the vocal tract, hold promise for applications in flexible and expressive speech synthesis. This kind of synthesis is typically called *articulatory synthesis*. Shadle and Damper (2002) outlined several complementary advantages that articulatory synthesizers should have over now widely adopted data-driven approaches like concatenative synthesis (Black, 2002) and Hidden Markov Model-based synthesis (Schroeter, 2006). Among these advantages are (a) the promise of producing speech associated with extraordinary speakers (e.g., an exceptional opera singer) or hypothetical speakers, from whom data can be difficult or impossible to collect, (b) the promise of changing the quality or type of speaker without having to perform additional statistical training of the synthesizer, (c) the promise of having meaningful parameters that can be helpful in fixing or adjusting the synthesizer output, in addition to providing insights into human speech production.

The models discussed here, in addition to being formal and mechanistic, are also causal, by intention of their development and by virtue of their historical context. Causal models can, as such, serve to encapsulate current theoretical understanding of the mechanisms underlying speech motor control into a compact and rigorous form. Analysis of speech behavior, even in response to challenging or contrived situations, may not always be sufficient for inferring the causal mechanisms of those behaviors. An individual’s sensorimotor behavior is, in general, the result of a complex mixture of stable and mature control mechanisms, learned and adaptive strategies, and possible individual-specific speaking strategies and impairments. Therefore, inferring the underlying mechanisms that contribute to observed behaviors is exceedingly difficult without an underlying framework. Neurologically relevant, mechanistic models of sensorimotor control provide a neurocomputational substrate which can aid in establishing causal relationships among the many component pathways and model parameters. By modeling and resynthesizing human behaviors, mechanistic models can infer the mechanisms underlying observed responses, including both impairment mechanisms and neural adaptation to those impairments. This process is termed *system identification* in an engineering context, and recent advances in methods for system identification have facilitated application to biological multivariate, closed-loop control systems (Engelhart *et al.*, 2016) and human sensorimotor control systems (Boonstra *et al.*, 2013; Engelhart *et al.*, 2015). Inroads have also recently been made in applying similar approaches in the domain of typical (Mitra *et al.*, 2010) and pathological (Ciccarelli *et al.*, 2016) speech motor control.

## V. CONCLUSION

In scanning the published literature on formal models of speech motor control, it is perhaps understandable to be left with the impression that a dizzying variety of qualitatively distinct models have been presented. Among all the models, DIVA and TD stand out as having a relatively long history of representation in the literature, and the efforts to develop them have remained almost entirely separate. SFC and FACTS make clear and related modeling contributions that enable the expressive power to explain specific empirical results in speech production. ACT is inspired by both DIVA and TD, but has a structure all its own. GEPPETO is the result of yet another distinct effort at model development; it is concerned with biomechanical considerations in the plant. Clearly, there is a healthy amount of variety in the various model architectures, especially in their specific use and method of combining the three essential functional components: feedforward, feedback and model predictive. However, it is nonetheless possible to view these models as belonging to a single, coherent framework. The present paper has attempted to cut through the difficulties associated with varying presentation and terminology, and to directly compare the models against the backdrop of such a framework. By presenting a clear comparison of the points of agreement and disagreement among the various models, as well as establishing areas where all models can be improved, this work can provide a foundation for future model development to improve our understanding of the speech motor system.

## RESOURCES

Several of the models discussed in this paper (DIVA, TD, CASY and the Maeda model) have been implemented as software tools, and are available for download online. Their addresses on the World Wide Web are included in the references below.

## ACKNOWLEDGMENTS

This work is sponsored by the Assistant Secretary of Defense for Research & Engineering (ASD[R&E]) under Air Force contract #FA8721-05-C-0002. The authors would like to thank Frank Guenther, Satrajit Ghosh and Bernd Kröger for their time and assistance in understanding the details of their models, as well as the three thoughtful reviewers of the manuscript.

## Appendix A

To aid the speech motor control practitioner, this Appendix consolidates the key algorithmic steps of three control architectures: Task Dynamics (TD), Directions into Velocities of Articulators (DIVA), and State Feedback Control (SFC). Bold lower case letters represent vectors, and bold upper case letters represent matrices. A single overhead dot represents a time derivative, and a double dot represents a second order time derivative.

## 1. Directions Into Velocities of Articulators (DIVA)

The Directions Into Velocities of Articulators (DIVA) model is a control architecture developed by (Guenther *et al.*, 2006) that uses a hybrid of feedback control and model predictive control. The model has been realized in software, and is available online (Nieto-Castanon, 2016).

## Algorithm

In the DIVA model predictive controller, the mobility space, **u**, and state of the plant, **x**, are identical, so **u** = **x**. Table II describes the variables in DIVA.

1. Compute a model-predictive control signal (termed *feedforward* in the published literature on DIVA).

a. Compute an error using the reference target in mobility space and the current predicted state of the plant.

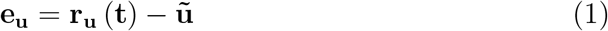
b. Compute a feedforward control update by scaling the error signal.

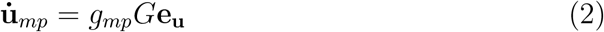
2. Compute a feedback-driven control signal using the reference target and the sensed plant output to compute an error in task space. Then, use a pseudoinverse Jacobian to convert the error from task space to mobility space. Do this in both the auditory and somatosensory feedback pathways.

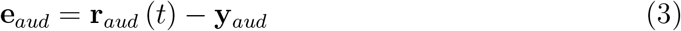

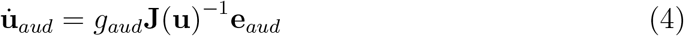

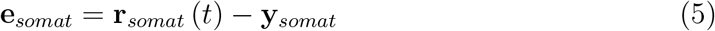

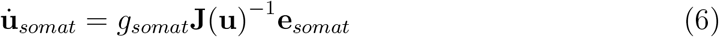
3. Combine the feedforward and feedback control updates to determine the new plant state.

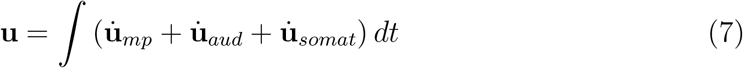

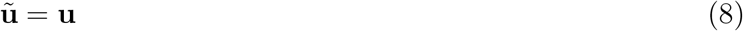

**TABLE II.** DIVA variables.

| Variable | Description |
| --- | --- |
| $\mathbf{e}_u$ | Error between reference target in mobility space and last command issued to the plant |
| $\mathbf{e}_{aud}, \mathbf{e}_{somat}$ | Error between the reference target in task space and sensed task space output |
| $\mathbf{r}_u(t)$ | Reference target in mobility space. Defined at each point in time as a region with a center and bounds of acceptable performance. |
| $\mathbf{r}_{aud}(t), \mathbf{r}_{somat}(t)$ | Reference target in task space. Defined at each point in time as a region with a center and bounds of acceptable performance. |
| $\dot{\mathbf{u}}_{aud}, \dot{\mathbf{u}}_{somat}$ | Change in mobility space position based on error in task space. A task space velocity update. |
| $\dot{\mathbf{u}}_{ff}$ | Change in mobility space position based on error in mobility space. A task space velocity update. |
| $\mathbf{u}$ | Mobility space position. Computed by integrating the feedforward and feedback mobility space velocities. |
| $\mathbf{y}_{aud}, \mathbf{y}_{somat}$ | Task space output |
| $g_{ff}$ | Gain applied to feedforward velocity update |
| $g_{aud}, g_{somat}$ | Gain applied to feedback velocity update |
| $G$ | Gain with a value between 0 and 1 that constrains velocities in mobility space from 0 to their maximum. |
| $\mathbf{J}(\mathbf{u})^{-1}$ | Pseudoinverse of the Jacobian. The pseudoinverse converts errors in task space to changes in velocity in mobility space. The pseudoinverse can be computed as the Moore-Penrose pseudoinverse. |

## 2. Task Dynamics

Task Dynamics is a feedback control architecture developed by (Saltzman and Kelso, 1987; Saltzman and Munhall, 1989). The architecture has been realized in software in the Task Dynamics Application (TADA) (Nam *et al.*, 2006) and available online (Nam, 2012).

## a. Algorithm

The Task Dynamics algorithm is described below, and all variables are defined in Table III.

1. Compute error in task space. In Task Dynamics, the task space, **y**, and the state, **x**, are identical, so **y** = **x**, and the error is

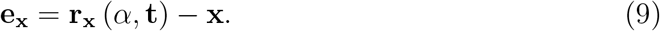
2. Use a dynamical system description of the controller, a second order ordinary differential equation, to compute the new acceleration state of the plant in task space as

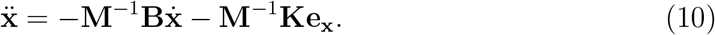
3. Use a pseudoinverse Jacobian to convert the task space acceleration to mobility space acceleration by

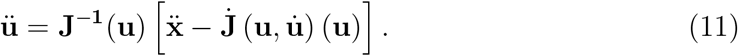
4. Integrate mobility space acceleration to get velocity and position in mobility space, so

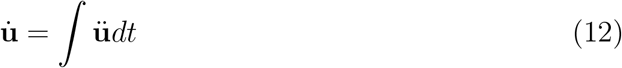

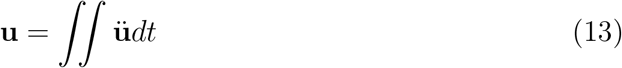

**TABLE III.** Task dynamic variables.

| Variable | Description |
| --- | --- |
| $\mathbf{x}, \dot{\mathbf{x}}, \ddot{\mathbf{x}}$ | Task space position, velocity, and acceleration, m by 1 vectors |
| $\mathbf{u}, \dot{\mathbf{u}}, \ddot{\mathbf{u}}$ | Mobility space position, velocity, and acceleration, n by 1 vectors |
| $\mathbf{r}_{\mathbf{x}}(\alpha)$ | Reference target in task space, m by 1 vector |
| $\mathbf{M}$ | Inertial coefficients, m by m diagonal matrix |
| $\mathbf{B}$ | Damping coefficients, m by m diagonal matrix |
| $\mathbf{J}$ | Jacobian transformation from mobility space to task space. An m by n matrix with elements $J_{ij} = \frac{\partial x_i}{\partial u_j}$ |
| $\mathbf{J}^{-1}$ | The (pseudo) inverse of the Jacobian. The Moore-Penrose pseudoinverse may be used, or other constraints can be applied to allow inversion of a non-square Jacobian. |
| $\dot{\mathbf{J}}$ | The time derivative of each element of the Jacobian. |

## 3. State Feedback Control

The State Feedback Control is a hybrid feedback/model-predictive control architecture proposed by (Houde and Nagarajan, 2011). Note that the notation used here follows the originally-published notation, and differs slightly from the simplified notation used in the main body of the present paper.

## a. Algorithm

1. Create a control update using the current estimate of the plant state by

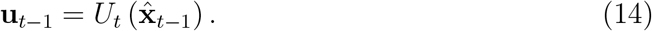
2. Create the new, true plant state using the true plant dynamics, *G*_*dyn*_, by

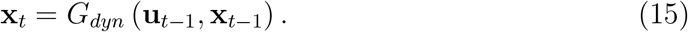
3. Create a new, predicted estimate of the plant state using the previous estimate of the plant state, 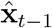, the previous control signal, **u**_*t*−1_, and an estimate of the plant dynamics, 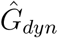, by

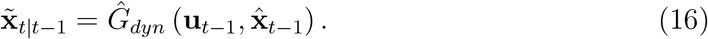

x
4. Generate the subsequent plant output using the true plant transformation from plant state to plant output by

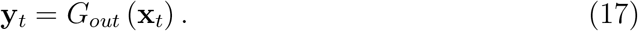
5. Create a correction term to the plant state estimate using the sensed feedback from the true plant by

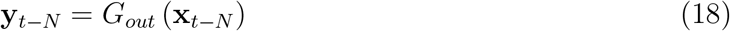

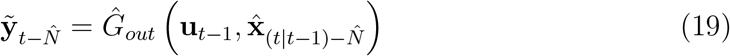

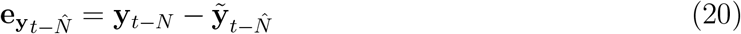

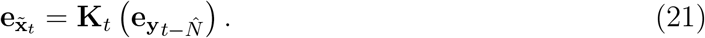
6. Combine the initial plant state estimate and the correction term to create the current estimate of the plant state by

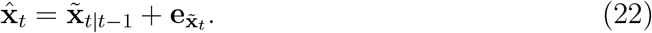

**TABLE IV.**
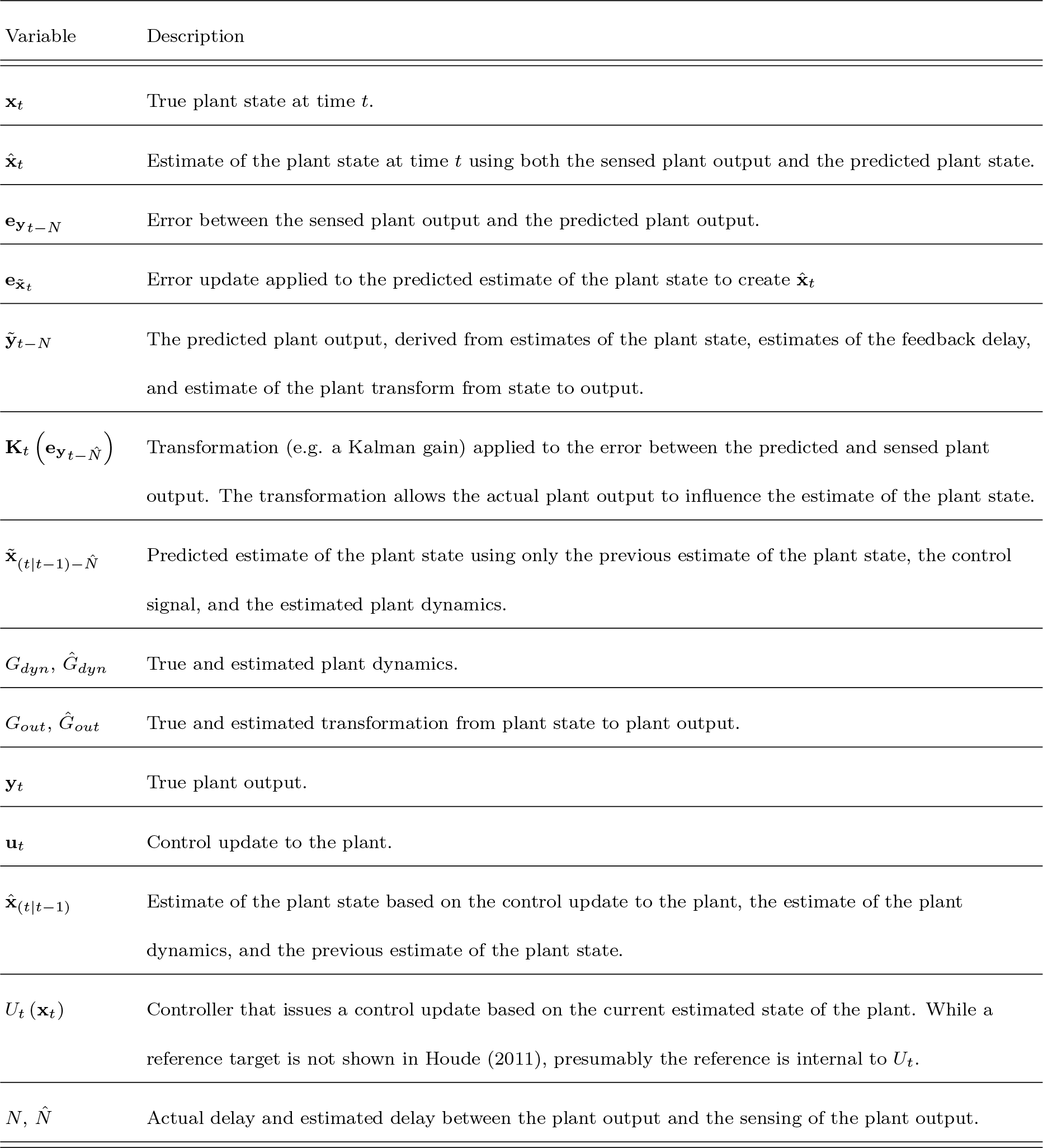
State feedback control variables.

## Appendix B

This appendix presents two articulatory speech synthesizers commonly referenced in the literature: the Configurable Articulatory Synthesizer (CASY), and the Maeda model. Bold lower case letters represent vectors, and bold upper case letters represent matrices. A single overhead dot represents a time derivative, and a double dot represents a second order time derivative.

## 4. Configurable Articulatory Synthesizer

The Configurable Articulatory Synthesizer (CASY) is a geometric model of the vocal tract based on the work of Mermelstein (1973) and developed by Rubin *et al.* (1996) and Iskarous *et al.* (2003). The governing equations are presented below, taken from Lammert (2013). The “q” variables in Lammert *et al.* (2013), that represent the articulators in mobility space, have been renamed to “u” in this paper for consistency of notation (see Tables V and VI for details about the variables/constants).

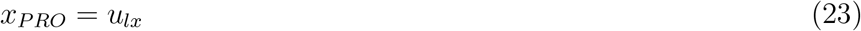

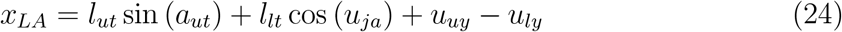

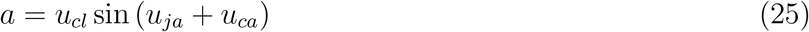

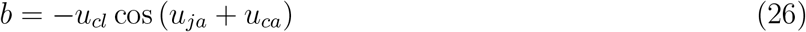

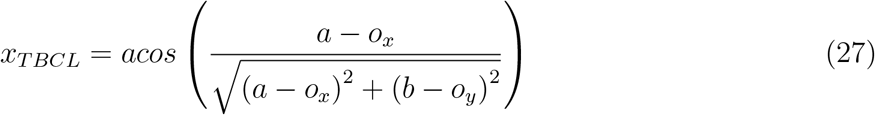

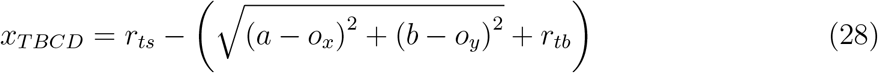

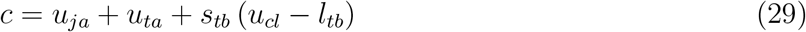

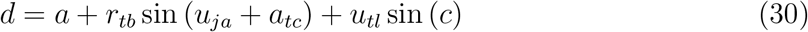

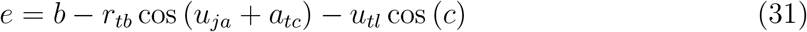

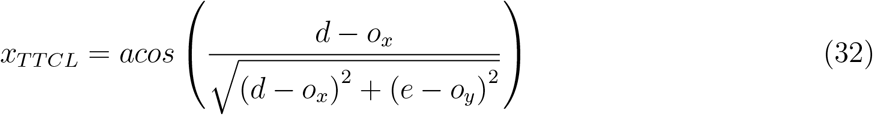

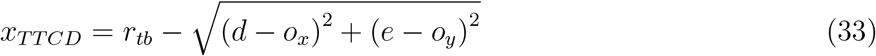

## 5. Maeda Articulatory Synthesizer

The Maeda articulatory speech synthesizer is a variable cross-sectional area, tube model of the vocal tract. Resonances of the tube can be computed, and these resonances are the formants. The formants can then be used to shape a vocal source (voiced or unvoiced) to create speech. A MATLAB instantiation of the Maeda synthesizer was created by Ghosh and available for download (Nieto-Castano, 2017).

**TABLE V.** CASY variables.

| Variable | Description |
| --- | --- |
| $x$ | Task space variable |
| $u$ | Mobility space variable |
| LX | Lip protrusion |
| UY | Upper lip vertical displacement |
| UT | Upper teeth |
| LY | Lower lip vertical displacement |
| JA | Jaw angle |
| CA | Tongue body angle |
| CL | Tongue body length |
| TL | Tongue tip length |
| TA | Tongue tip angle |
| LA | Lip aperture |
| PRO | Lip protrusion |
| TBCD | Tongue body constriction degree |
| TBCL | Tongue body constriction location |
| TTCD | Tongue tip constriction degree |
| TTCL | Tongue tip constriction location |

Ciccarelli (Ciccarelli, 2017) created a polynomial approximation to the vocal tract component to allow fast formant computation and fast, tractable computation of the pseudoinverse of the Jacobian. The polynomial approximation was determined by running the Ghosh implementation of the Maeda model across a set of parameters, uniformly sampled from the mobility space of the model, to create a lookup table of parameters and formant values.

**TABLE VI.**
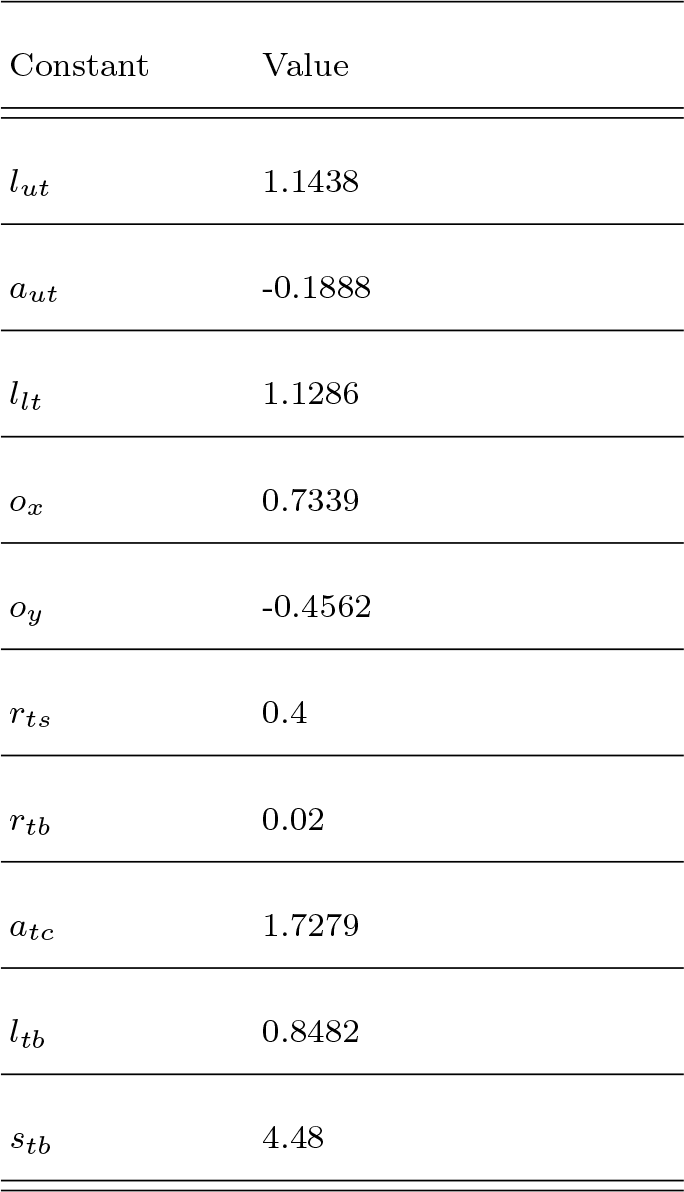
CASY constants.

Formant points outside the standard vowel quadrilateral as determined by visual inspection were excluded. The remaining pairs of articulator points and formants were then fit using a least squares polynomial approximation. The order of the polynomial was a compromise between the fit to the data and the complexity of the polynomial. It was found that a second order polynomial achieved a reasonable balance between these two requirements. While the mapping from articulators to formants is preserved to within a certain error, it has not been evaluated whether the relationship between articulators encoded by the polynomial fundamental alters the trajectories of articulators in previous implementations of the Maeda model.

## Footnotes

1 In the speech motor control literature, the term ‘articulatory space’ is often used instead of ‘mobility space’. The latter term is adopted from the robotics literature (Sciavicco *et al.*, 2012) here to provide a neutral terminology for referring specifically to the configuration of the plant, whereas terminology used in the literature often leads to confusion over whether the term ‘articulatory’ refers to low-level descriptions of the plant or high-level tasks spaces defined in articulatory terms.

2 For this example, the simplifying assumption is made that the feedback signal is in task space, i.e. **y**_**x**_

3 Optimal here means closest to the true state of the plant, where “closest” means having the smallest mean squared error

4 The description of SFC presented here uses a different notation than in Houde and Nagarajan (2011), simplified for clarity of presentation. For a more complete mathematical description, see Appendix A.

